# Diminished ascending drive in transdiagnostic anhedonia shifts the cortico-subcortical hierarchy away from bottom-up dominance

**DOI:** 10.64898/2026.10.12.758297

**Authors:** Payman Roghani, Jakub Vohryzek, Morten L. Kringelbach, Gustavo Deco

## Abstract

Anhedonia, the diminished capacity to experience or pursue pleasure, is a core feature of mood and psychotic disorders, yet the large-scale brain dynamics underlying it remain uncharacterized. Dominant accounts posit striatal hypofunction with compensatory prefrontal over-recruitment, but they rest on activation magnitudes and symmetric correlations, which cannot resolve the direction of influence. Here, in 210 adults from the Transdiagnostic Connectome Project, we defined symptom-derived low- and high-anhedonia subgroups and estimated each participant’s directed effective connectivity using a whole-brain generative model fitted to resting-state fMRI. In high anhedonia, the cortex shifted upstream relative to the subcortex, flattening the cortico-subcortical hierarchy. Correspondingly, the net balance of direct cortico-subcortical flow shifted from bottom-up to balanced. This shift recurred within the frontostriatal reward circuit and replicated across independent halves of each participant’s data. Flow asymmetry also increased with subject-level severity. Decomposing flow into ascending and descending arms showed that both weakened in high anhedonia, but ascending subcortex-to-cortex and striatum-to-prefrontal drive fell more than the descending drive. These findings identify a reorganization of cortico-subcortical hierarchy in anhedonia and suggest that the previously reported prefrontal over-recruitment reflects diminished ascending drive rather than stronger cortical output.

## Introduction

Anhedonia, the loss of pleasure or of the motivation to pursue it, is one of the two cardinal symptoms of major depressive disorder (MDD) and a transdiagnostic feature of mood, anxiety, and psychotic illness^1–4^. It predicts a chronic course, poor treatment response, and elevated suicide risk^5^, yet it remains a major unmet therapeutic need^3,6,7^. Because anhedonia cuts across diagnostic boundaries and is only partially captured by any single disorder, it is a prototypical target for the dimensional, neurobiologically grounded approach advocated by the Research Domain Criteria (RDoC) framework, which seeks mechanisms that span Diagnostic and Statistical Manual of Mental Disorders (DSM) categories^8^. Consistent with this, transdiagnostic studies find reward-circuit dysfunction that tracks anhedonia severity independent of diagnosis^9–11^.

The neurobiology of anhedonia centers on the mesocorticolimbic reward circuit, in which dopaminergic projections from the midbrain reach the ventral and dorsal striatum and the prefrontal cortex (PFC), including orbitofrontal, ventromedial, and anterior cingulate regions^3,12–14^. In functional MRI (fMRI) studies, both during reward tasks and at rest, anhedonia has been linked to blunted striatal responses accompanied by prefrontal hyperactivation, an imbalance widely interpreted as compensatory top-down control over a hyporesponsive striatum^3,15–18^. This influential account, however, rests largely on regional activation magnitudes and on symmetric, undirected functional connectivity, neither of which can specify the direction of influence between PFC and striatum. Whether anhedonia is accompanied by a genuine reorganization of directed cortico-subcortical and frontostriatal information flow therefore remains untested.

Addressing this requires a measure that is both directed and grounded in brain dynamics. Static functional connectivity, the statistical relationship between the blood-oxygen-level-dependent (BOLD) time series of two regions and conventionally quantified as the Pearson correlation between them, is symmetric by construction and therefore blind to hierarchy of regional interactions, while regional activation does not describe how regions influence one another.

Whole-brain generative modeling overcomes both limitations: by fitting a dynamical model to empirical resting-state activity and iteratively adjusting the strengths of anatomically supported connections, it yields a subject-specific, directed connectome, the generative effective connectivity (GEC), that offers a mechanistic account of the data^19–22^. Directionality is recoverable because the fit is constrained not only by the symmetric functional connectivity but also by the time-shifted (lagged) covariance between regional signals, whose inherent asymmetry breaks the symmetry of the coupling and assigns a direction of influence that static connectivity cannot resolve^20,23^. Applied to the GEC, graph-theoretic measures of trophic level and trophic coherence, adapted from the analysis of directed networks in ecology, place each region within the brain’s directed hierarchy and summarize how strongly that hierarchy is expressed^24^. Trophic levels capture each region’s position in the global hierarchy, integrating both direct and indirect pathways. We pair them with a complementary, mathematically related read-out of the GEC matrix: the net balance of directed flow across a chosen boundary between two compartments, computed only from the connections that cross it. Its added value is that the flow in each direction can be tested separately, distinguishing a change in top-down drive from a change in bottom-up drive. These hierarchy read-outs have proven more sensitive than conventional connectivity-based measures for distinguishing brain states^20,25^.

Here we apply this framework to transdiagnostic anhedonia in the Transdiagnostic Connectome Project (TCP)^26^, a richly phenotyped open dataset spanning healthy controls and individuals across affective, anxiety, and psychotic disorders. In this cohort, Cocuzza et al.^27^ recovered clinically relevant variance from intrinsic and task-evoked functional connectivity, classifying primary diagnosis and multi-dimensional symptom profiles above chance. Their measures were undirected, however, and whether this variance has a directional, hierarchical organization has not been tested. Here, we defined dimensional low- and high-anhedonia subgroups from a multi-scale factor analysis of three self-report pleasure and reward scales: the Snaith–Hamilton Pleasure Scale (SHAPS)^28^, the Temporal Experience of Pleasure Scale (TEPS)^29^, and the Depression Anxiety Stress Scale (DASS)^30^. We then estimated each participant’s resting-state GEC. We further asked whether and how the brain’s directed hierarchy differs with anhedonia and whether any such difference scales with severity and exceeds what matched undirected measures detect. If the compensatory-control account is correct, high anhedonia should be marked by a prefrontal move toward a more upstream position relative to the striatum and a shift of cortico-subcortical and frontostriatal flow in the top-down direction. Whether such a shift reflects heightened prefrontal output or a loss of ascending drive can only be resolved by decomposing the directed flow into ascending and descending arms, a distinction invisible to activation and undirected measures.

## Results

The analysis proceeded in three steps (**Figure 1**). We first derived low- and high-anhedonia subgroups by submitting items from the three pleasure–reward scales to a polychoric exploratory factor analysis and clustering the resulting factor scores with a Gaussian mixture model (**Figure 1a**). For every participant we then fitted a whole-brain Hopf model to resting-state fMRI from 232 cortical and subcortical regions, seeded with structural connectivity and constrained by both the functional connectivity and the time-shifted covariance, to obtain a directed GEC matrix (**Figure 1d**; Methods).

**Figure 1.**
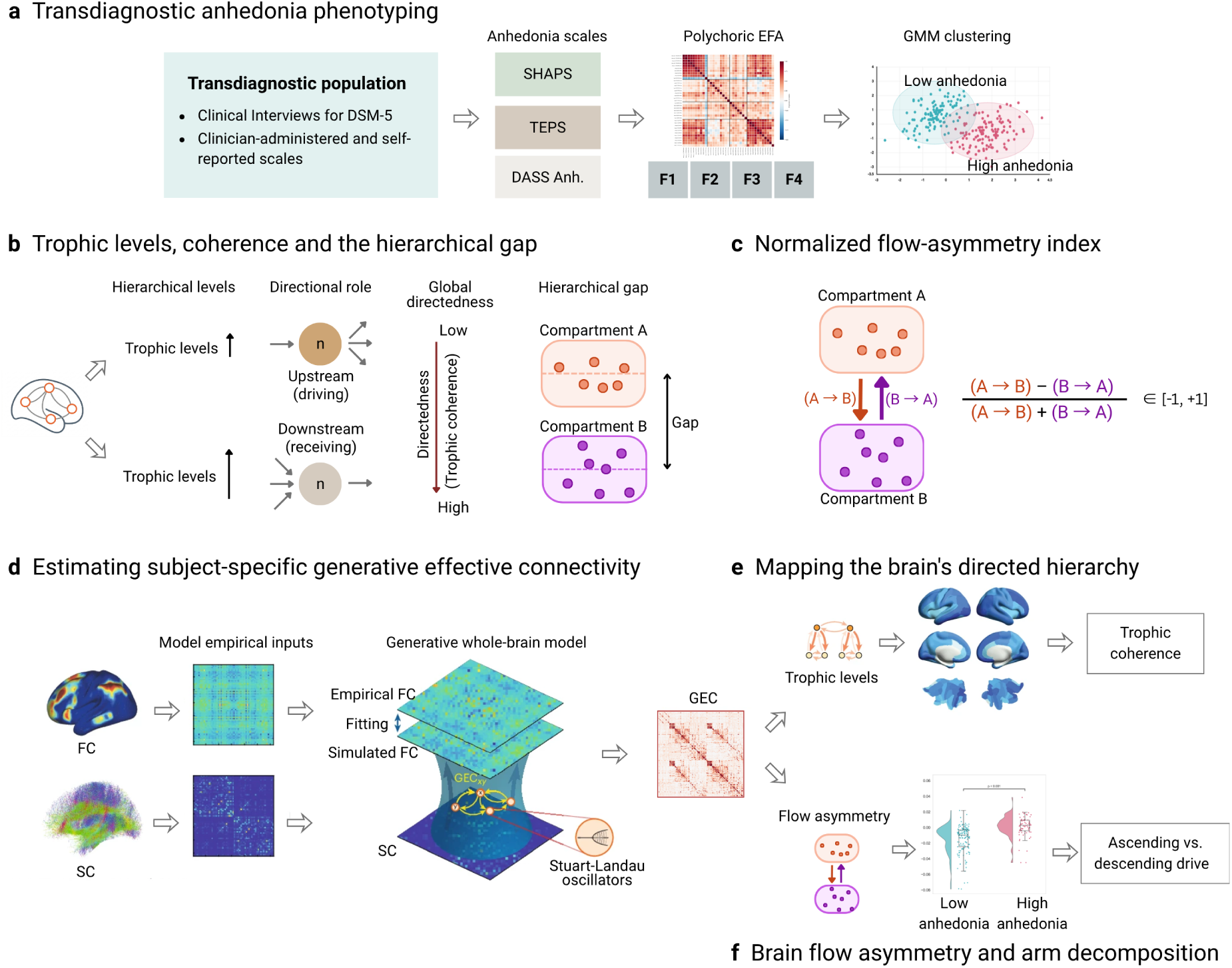
Dataset and analysis-pipeline overview. **(a)** Items from three pleasure–reward scales (SHAPS, TEPS, DASS) enter a polychoric exploratory factor analysis; a Gaussian mixture model on the resulting factor scores partitions the transdiagnostic sample into low- and high-anhedonia subgroups. **(b)** A region’s trophic level reflects its position in the brain’s directed hierarchy, determined by the imbalance of incoming and outgoing connection weights. Regions whose outgoing weight exceeds their incoming weight have lower trophic levels and occupy an upstream (driving) position, whereas regions where this balance is reversed have higher trophic levels and occupy a downstream (receiving) position. Trophic coherence summarizes the whole network, quantifying how consistently edges align along this hierarchy. The hierarchical gap is the difference in mean trophic level between two compartments (dashed lines), indexing how far one sits upstream of the other. **(c)** A normalized index contrasts directed flow from compartment A→B (the descending arm) against B→A (the ascending arm), bounded in [−1, +1], capturing which compartment exerts the greater directed influence. Because the two arms are summed separately, each can also be tested on its own as a measure of descending and ascending drive, attributing a shift in the index to a change in either arm. **(d)** A Hopf model of coupled Stuart–Landau oscillators, seeded with structural connectivity and fitted to each subject’s empirical functional connectivity and its time-shifted covariance, yields a directed generative effective connectivity (GEC) matrix. **(e)** Trophic levels computed on the GEC place each region and compartment within the brain’s directed hierarchy and summarize its overall directedness (trophic coherence). **(f)** The flow-asymmetry index is evaluated across the cortico-subcortical (and frontostriatal) boundary and compared between subgroups; its two arms are then tested separately to attribute any group shift to a change in ascending or descending drive. EFA, exploratory factor analysis; GMM, Gaussian mixture model; FC, functional connectivity; SC, structural connectivity.

From each GEC matrix we derived two complementary read-outs. The first is trophic level^20,24^, which places each region in the directed hierarchy according to the balance between the GEC weight it sends and receives, integrating indirect, multi-hop pathways as well as direct connections. Regions that send more than they receive have lower trophic levels and sit upstream; regions that receive more sit downstream. Averaging within a compartment gives its mean position, and the difference between two compartment means (the hierarchical gap) indicates how far downstream one lies relative to the other. Trophic coherence is the global summary, with higher values indicating a more strongly directed organization and lower values a flatter one (**Figure 1b,e**).

The second read-out is a flow-asymmetry index: the total GEC weight crossing a boundary (e.g. cortex to subcortex) in one direction against the total in the other, normalized to [−1, +1] so that it reflects the balance rather than the magnitude of flow (**Figure 1c,f**). It is computed only from the connections that cross the boundary, and its added value is that its two directional sums can be tested separately as descending and ascending arms.

In our analysis we moved from global trophic coherence to regional and compartmental trophic levels, and then to the balance of flow across the cortico-subcortical and frontostriatal boundaries and its two arms, asking throughout whether effects localized to the reward circuit. Group differences were tested with ordinary least squares (OLS) regression in three models (full sample adjusted for age, sex, and patient/healthy-control status; patients only; patients only adjusted for broad diagnostic category), with Benjamini–Hochberg false discovery rate (FDR) correction within each family of related metrics. Flow-asymmetry measures were also correlated with symptom severity, and the key analyses were repeated on undirected functional connectivity.

### Transdiagnostic anhedonia subgroups

We first defined anhedonia dimensionally and transdiagnostically, drawing items from three self-report scales in the TCP behavioral battery: the SHAPS, the TEPS, and a five-item anhedonia composite from the DASS. The SHAPS and TEPS are dedicated measures of hedonic capacity, while the DASS composite captures anhedonic features within a broader depression–anxiety–stress inventory. Parallel analysis supported a four-factor solution (Methods). Rather than simply reproducing the three scales, the four factors cut across them: items from each instrument loaded on more than one factor, the TEPS split into separate anticipatory and consummatory factors, and the DASS items formed their own distress-related factor (**Figure 2a**). Because the underlying dimensions did not line up with any single questionnaire, combining items from all three scales captured more of this structure than any one scale with a fixed cut-off could.

**Figure 2.**
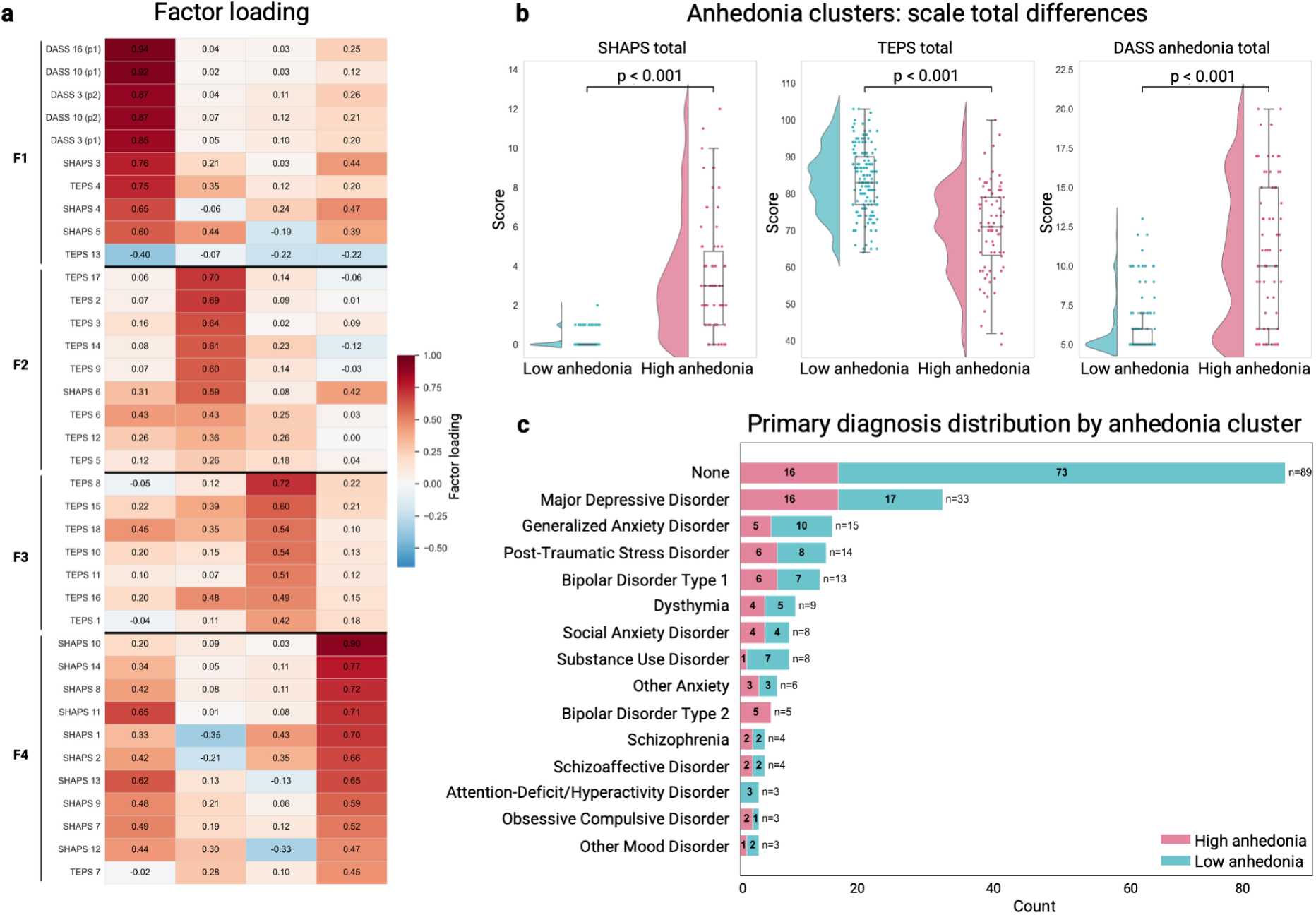
A multi-scale factor solution defines transdiagnostic low- and high-anhedonia subgroups. **(a)** Factor-loading matrix from a polychoric exploratory factor analysis of the SHAPS, TEPS, and DASS anhedonia items, showing that the four factors cut across the three source scales. Rows are items, grouped by the factor they load on most strongly (F1–F4); cell values are factor loadings, with colour encoding their sign and magnitude (colour bar). **(b)** The low- and high-anhedonia subgroups separate cleanly on all three scale totals (all FDR-corrected p < 0.001). Each panel shows a half violin (kernel density, normalized to equal width), a box (median and interquartile range, whiskers 1.5 × interquartile range, outliers omitted) and one dot per participant (low anhedonia n = 145; high anhedonia n = 74). **(c)** Cluster membership by primary diagnosis: high anhedonia spans nearly every category but is enriched in mood and psychotic disorders. Bars give the number of participants per subgroup, with the total per diagnosis at the right.

Fitting a Gaussian mixture model to the four factor scores yielded a low-anhedonia subgroup (n = 145) and a high-anhedonia subgroup (n = 74). The solution separated cleanly on all three scale totals, with large effect sizes (SHAPS 3.35 ± 3.07 vs. 0.15 ± 0.37, Cohen’s d = 1.78; DASS anhedonia 2.15 ± 0.92 vs. 1.19 ± 0.36, d = 1.58; TEPS 3.08 ± 0.69 vs. 2.36 ± 0.50, d = 1.26; all FDR-corrected p < 0.001; **Figure 2b**). High anhedonia appeared across nearly every diagnostic category in the sample, but was strongly enriched in mood and psychotic disorders (e.g. MDD 48.5%, bipolar I 46.2%, schizophrenia 50.0%) relative to participants with no primary diagnosis (**Figure 2c**). The grouping is therefore best read as a dimensional, transdiagnostic phenotype that is over-represented in, but not confined to, mood and psychotic conditions.

Within the subset with complete imaging and demographic data (n = 210; low anhedonia n = 139, high anhedonia n = 71), the groups did not differ in age (medians 27.0 vs. 31.0 years; means 31.7 ± 11.8 vs. 35.5 ± 15.2; Wilcoxon rank-sum Z = −0.84, p = 0.40) or sex (χ²(1) = 2.73, p = 0.099) but were strongly imbalanced on clinical status: only 19.7% of the high-anhedonia group were healthy controls versus 48.9% of the low-anhedonia group (χ²(1) = 15.63, p < 0.001), which is the reason clinical status is carried as a covariate throughout.

### Global hierarchical directedness is unchanged

We first asked whether anhedonia was associated with the overall strength of hierarchical directedness, measured by trophic coherence, and it was not. Whole-brain trophic coherence did not differ between groups in the covariate-adjusted model (β = 0.0003, 95% CI [−0.0014, 0.0020], p = 0.69), and the null held in the patients-only (p = 0.74) and diagnosis-adjusted (p = 0.71) models. Computing coherence separately within the cortical and subcortical subnetworks gave the same result (cortical p = 0.93; subcortical p = 0.81).

### Regional trophic levels: an upstream cortical shift

Regional trophic levels, unlike the global scalar, differed between groups. Across all 232 regions, 122 differed at an uncorrected threshold and 29 survived FDR correction (q < 0.05; **Figure 3a**).

**Figure 3.**
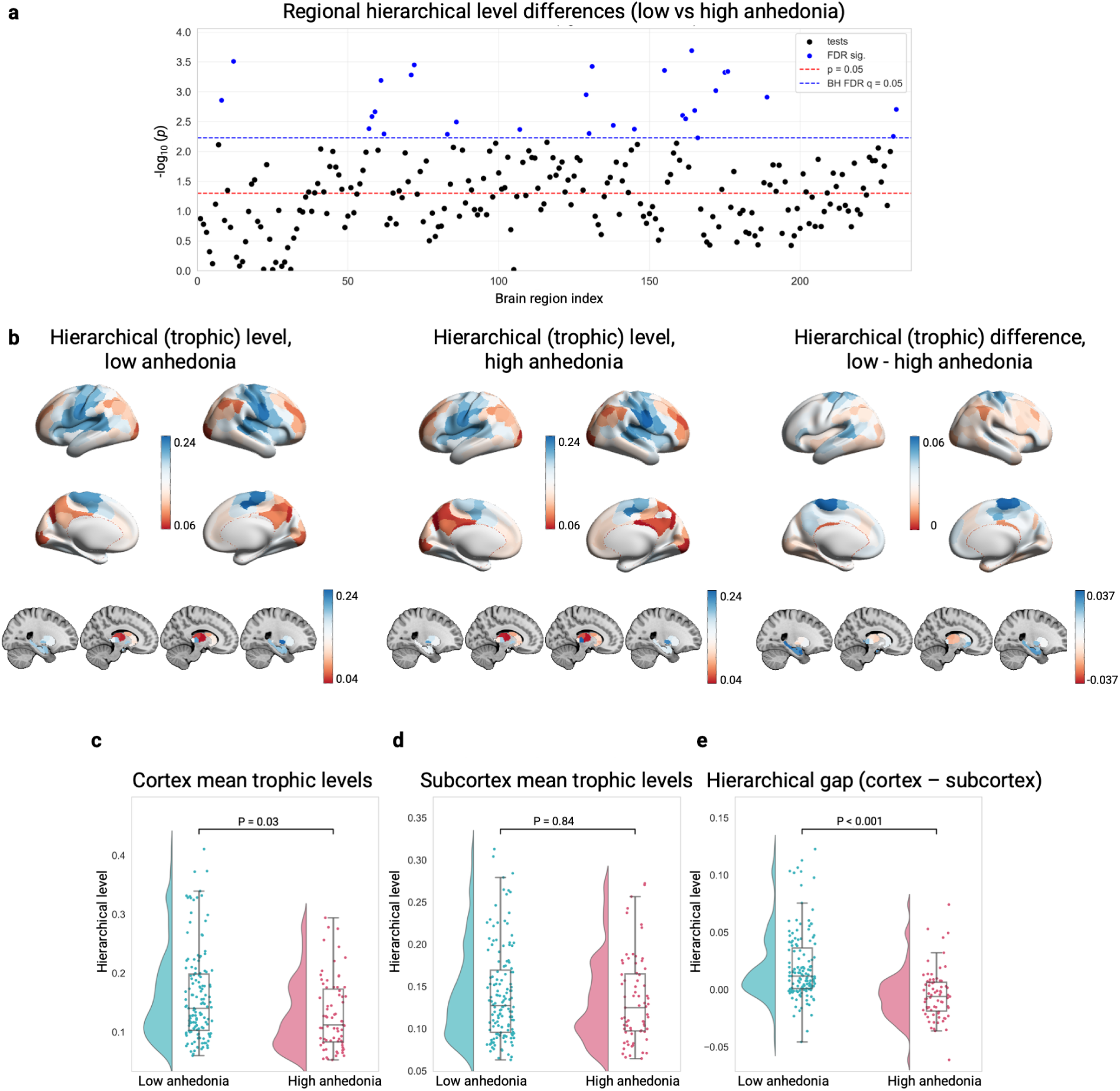
Cortex shifts upstream relative to subcortex, flattening the cortico-subcortical hierarchy in high anhedonia. **(a)** Regional trophic-level group differences across all 232 regions; one dot per region, plotted as −log10(p) against region index, with dashed lines marking the uncorrected p = 0.05 and Benjamini–Hochberg FDR q = 0.05 thresholds. Twenty-nine regions survive FDR correction, the great majority cortical and lower in high anhedonia, with two subcortical regions higher. **(b)** Cortical (surface) and subcortical (slices) trophic levels in the low- and high-anhedonia groups and their difference (low minus high), showing a broadly upstream cortical shift relative to each participant’s most upstream region; colour encodes trophic level, scaled as in the adjacent colour bars. **(c)** Mean cortical trophic level is lower in high anhedonia (p = 0.03), whereas **(d)** the subcortical mean does not differ (p = 0.84); both are relative to each participant’s most upstream region. **(e)** The cortex–subcortex hierarchical gap is robustly reduced in high anhedonia (p < 0.001), the strongest of the three measures. In **c**–**e**, each panel shows a half violin (kernel density, normalized to equal width), a box (median and interquartile range, whiskers 1.5 × interquartile range, outliers omitted) and one dot per participant (low anhedonia n = 139; high anhedonia n = 71).

These differences were strikingly systematic. Twenty-seven of the 29 surviving regions were cortical and all showed the same sign, with lower trophic levels in high anhedonia, indicating a position closer to each participant’s most upstream region (Methods). These regions were anatomically clustered: somatomotor cortex was most heavily represented (11 parcels), followed by posterior dorsal-attention (5), posterior default-mode (6), salience/ventral-attention (3), and right visual cortex (2). In contrast, only two subcortical regions survived correction – the right dorsoanterior thalamus and right anterior globus pallidus – and these had higher trophic levels in high anhedonia. This shift is thus overwhelmingly cortical (**Figure 3b**), with only these two subcortical exceptions.

### A reduced cortex–subcortex hierarchical gap in high anhedonia

Summarizing this pattern at the level of whole compartments, the mean cortical trophic level was significantly lower in high anhedonia (β = −0.0247, 95% CI [−0.0467, −0.0028], p = 0.027), whereas the mean subcortical level did not differ (β = −0.0017, p = 0.84; **Figure 3c,d**). Because these single-compartment means are measured relative to each participant’s most upstream region, which differs between participants, they do not by themselves show which compartment moved (Methods). The measure that does not depend on this reference is the cortex–subcortex hierarchical gap (the mean cortical minus the mean subcortical trophic level, indexing how far cortex sits downstream of subcortex in the hierarchy), which was robustly reduced in high anhedonia and was by far the strongest of the three compartment measures (β = −0.0230, 95% CI [−0.0316, −0.0144], p < 0.001; d = −0.83; R² = 0.169 vs. 0.081 and 0.044 for the single-compartment models; **Figure 3e**). The gap was positive in low anhedonia (mean = +0.020), with cortex downstream of subcortex, but close to zero in high anhedonia (mean = −0.005; covariate-adjusted mean −0.003, 95% CI [−0.010, +0.004], p = 0.36), eliminating the bottom-up ordering seen in low anhedonia. Although modest in raw units (about 8% of the average within-participant range of trophic levels), this shift was large relative to between-participant variability. This gap effect was the most reproducible finding in the study: it was unchanged or strengthened when restricted to patients (β = −0.0254, p < 0.001) and when broad diagnostic category was added (β = −0.0264, p < 0.001, R² = 0.303), demonstrating that this shift tracks anhedonia rather than clinical status or diagnosis.

### Network-level trophic levels

Averaging trophic levels within each of the seven canonical functional networks^31^ and within the subcortex, all seven cortical networks were lower in high anhedonia relative to each participant’s most upstream region (largest in somatomotor and default-mode cortex), whereas the subcortex did not differ, but none survived FDR correction across the eight comparisons (all q ≈ 0.07; Supplementary Fig. 1).

### The same cortex–subcortex shift recurs within the reward circuit

We next asked whether the brain-wide pattern, cortex shifting upstream relative to subcortex, recurred within the mesocorticolimbic reward circuit specifically – the network most directly implicated in anhedonia. We grouped regions into seven reward regions of interest (ROIs) spanning both sides of that boundary: four cortical components (orbitofrontal cortex (OFC), medial/ventromedial prefrontal cortex (mPFC/vmPFC), anterior cingulate cortex (ACC), and dorsolateral prefrontal cortex (dlPFC)) and three striatal targets (nucleus accumbens (NAc), caudate, putamen). The same pattern emerged (**Figure 4a**). All four cortical reward nodes had lower trophic levels in high anhedonia (OFC β = −0.0250; mPFC/vmPFC β = −0.0271; ACC β = −0.0232; dlPFC β = −0.0213), each significant uncorrected, whereas the three striatal nodes did not differ between groups (NAc β = −0.0089, p = 0.29; caudate β = +0.0035, p = 0.66; putamen β = +0.0052, p = 0.61); the dorsal striatal coefficients were small and positive, a non-significant trend opposite in sign to the cortex. The cortical effects strengthened as confounding variance was removed: marginal in the full sample (q = 0.062–0.081), they survived FDR correction in the patients-plus-diagnosis model (all four q = 0.045), while the striatal nodes remained non-significant throughout (Supplementary Table 1). Relative to each participant’s most upstream region, the cortical components of the reward circuit thus sit further upstream in high anhedonia, whereas the striatal nodes do not differ.

**Figure 4.**
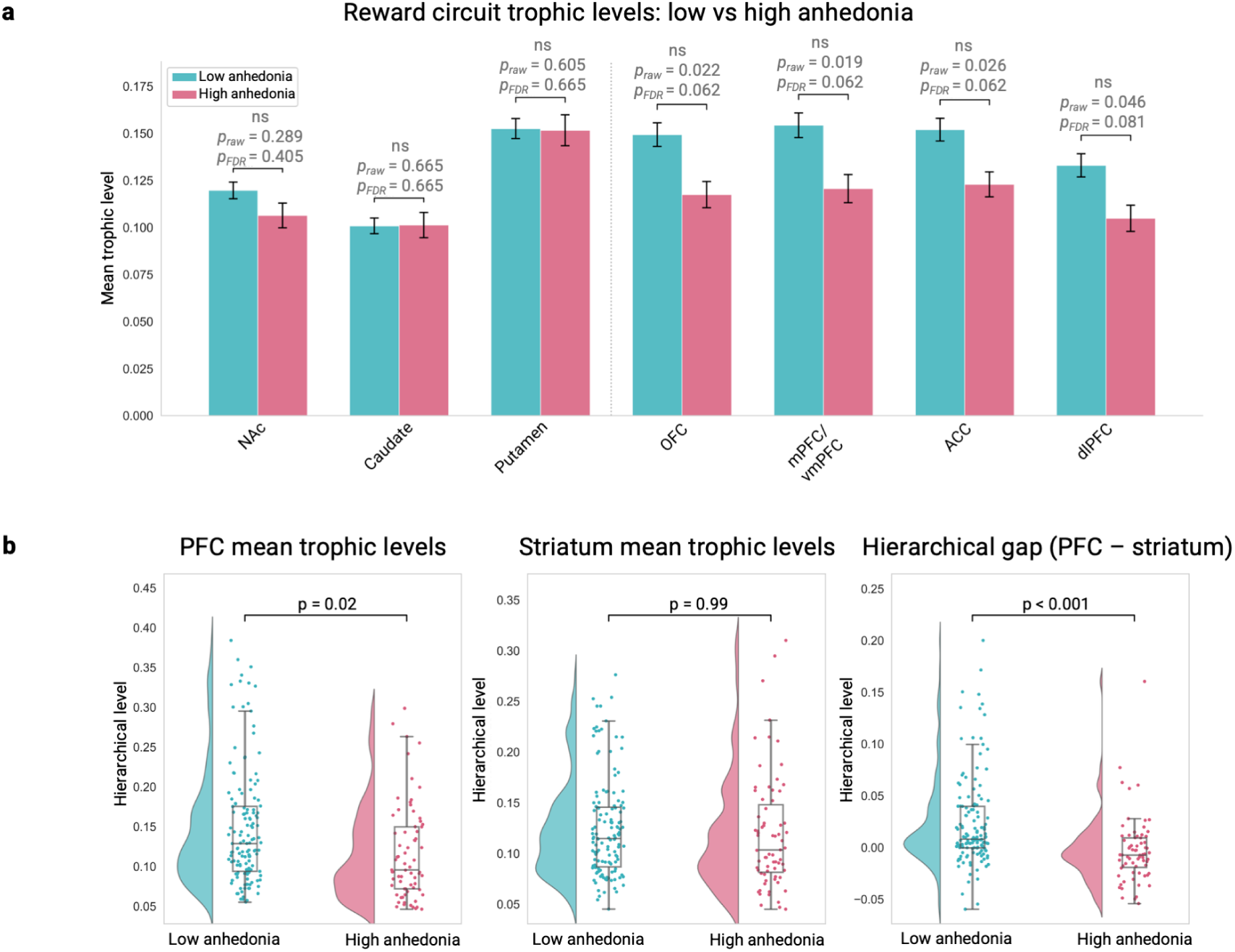
The same cortex–subcortex shift recurs within the reward circuit. **(a)** Mean trophic levels for the seven reward ROIs. All four cortical components (OFC, mPFC/vmPFC, ACC, dlPFC) are lower in high anhedonia (each significant uncorrected), whereas the three striatal nodes (NAc, caudate, putamen) do not differ; bars show group means, with error bars giving the standard error of the mean (SEM). Above each pair, p_raw is the uncorrected p value and p_FDR the value after Benjamini–Hochberg correction across the seven ROIs (ns, not significant after correction). **(b)** Collapsed to compartments, the mean prefrontal trophic level is lower in high anhedonia (p = 0.02) while the striatal mean does not differ (p = 0.99; both relative to each participant’s most upstream region), and the PFC–striatum hierarchical gap is robustly reduced (p < 0.001). Each panel shows a half violin (kernel density, normalized to equal width), a box (median and interquartile range, whiskers 1.5 × interquartile range, outliers omitted) and one dot per participant (low anhedonia n = 139; high anhedonia n = 71).

### A reduced frontostriatal hierarchical gap

Collapsing the reward nodes into a prefrontal compartment (OFC + mPFC/vmPFC + ACC + dlPFC) and a striatal compartment (NAc + caudate + putamen) reproduced the whole-brain gap result within the reward circuit. Mean PFC trophic level was lower in high anhedonia (β = −0.0246, 95% CI [−0.0460, −0.0033], p = 0.024), the striatal mean was essentially identical between groups (β = −0.00006, p = 0.99), and the PFC–striatum gap was therefore robustly reduced (β = −0.0246, 95% CI [−0.0371, −0.0120], p < 0.001; d = −0.60; R² = 0.103; **Figure 4b**), shifting from positive in low anhedonia (mean = +0.024) to close to zero in high anhedonia (mean = −0.003). As with the whole-brain gap, the frontostriatal gap was the strongest and most reproducible of the three measures and strengthened across the sensitivity models (patients only β = −0.0271, p < 0.001; patients + diagnosis β = −0.0281, p < 0.001).

### Directed flow shifts away from bottom-up dominance at the cortico-subcortical boundary

Our next analysis turned to the direct cortico-subcortical boundary. For each subject we summed the GEC weights running from cortex to subcortex (*A*) and, separately, from subcortex to cortex (*B*), and formed a normalized flow-asymmetry index, (*A* − *B*) / (*A* + *B*); positive values indicate net top-down (cortex-driving) flow and negative values net bottom-up flow (**Figure 1c**). The balance shifted significantly in the top-down direction in high anhedonia (**Figure 5a**): low-anhedonia subjects were on average net bottom-up (mean index = −0.0148), whereas high-anhedonia subjects were balanced (mean = +0.0014; covariate-adjusted mean +0.0005, 95% CI [−0.004, +0.005]), a robust group effect (β = +0.0150, 95% CI [+0.0099, +0.0201], p < 0.001; d = +0.90; R² = 0.205) that strengthened in the patients-only (β = +0.0162, p < 0.001) and diagnosis-adjusted (β = +0.0167, p < 0.001; R² = 0.328) models. The effect held on the raw, unnormalized difference, confirming it does not arise from the smaller total coupling in high anhedonia.

**Figure 5.**
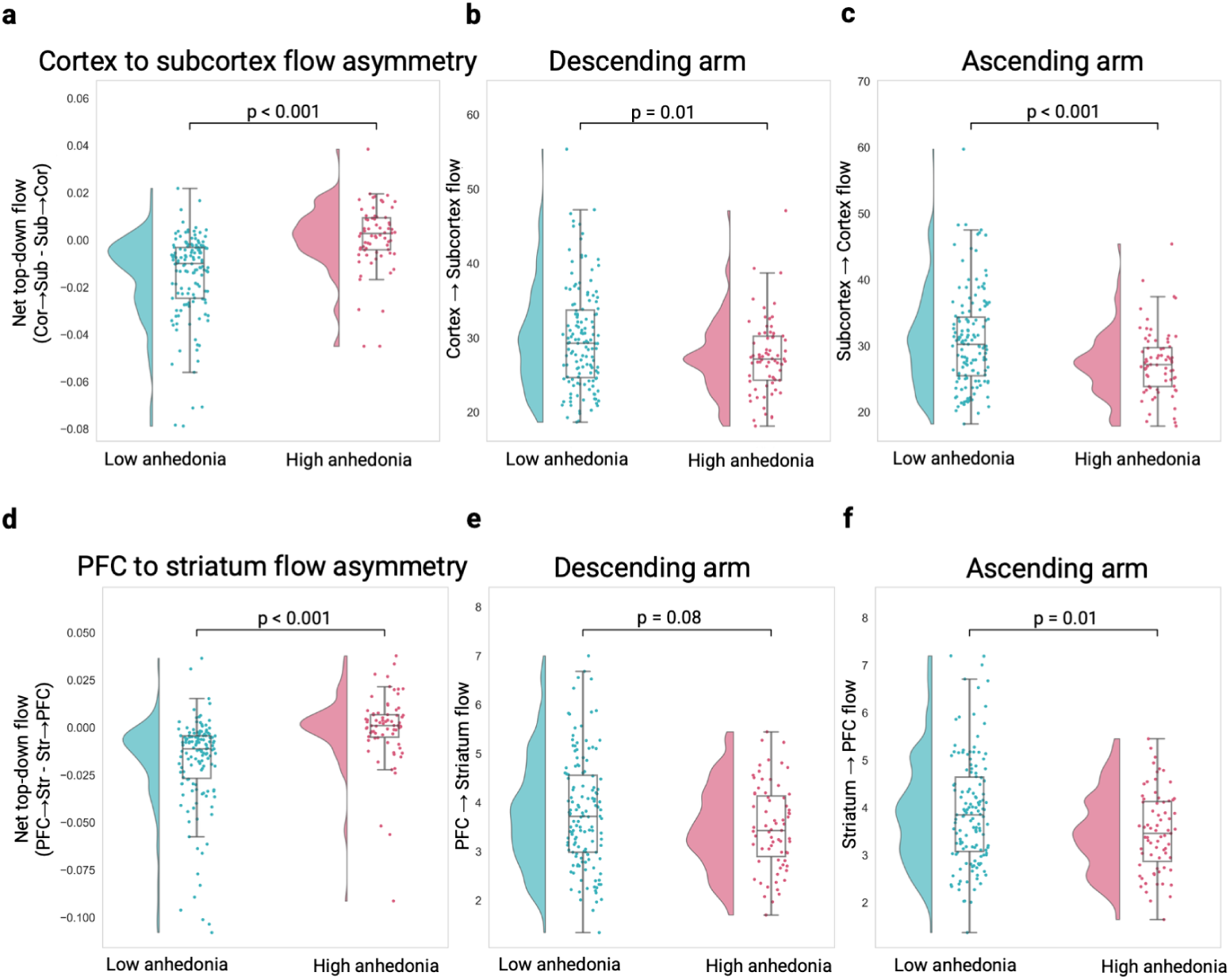
Directed flow shifts away from bottom-up dominance through a disproportionate loss of ascending drive. **(a)** The normalized cortex–subcortex flow-asymmetry index shifts in the top-down direction in high anhedonia (p < 0.001), from net bottom-up in the low group to balanced in the high group. **(b, c)** Decomposing the boundary into its two arms: both the descending cortex→subcortex arm (**b**, p = 0.01) and the ascending subcortex→cortex arm (**c**, p < 0.001) are weaker in high anhedonia, with the ascending arm falling more, which removes the net bottom-up balance. **(d)** The same top-down shift is reproduced within the reward circuit for the PFC–striatum asymmetry (p < 0.001). **(e, f)** Its arm decomposition mirrors the whole-boundary pattern: the descending PFC→striatum arm declines only weakly (**e**, p = 0.08) while the ascending striatum→PFC arm falls significantly (**f**, p = 0.01). Throughout, each panel shows a half violin (kernel density, normalized to equal width), a box (median and interquartile range, whiskers 1.5 × interquartile range, outliers omitted) and one dot per participant (low anhedonia n = 139; high anhedonia n = 71). Axis labels: Cor, cortex; Sub, subcortex; Str, striatum.

### The top-down shift is reproduced within the frontostriatal circuit

Applying the same analysis to the reward circuit specifically (PFC↔striatum) mirrored the whole-boundary finding (**Figure 5d**). Low-anhedonia subjects were net bottom-up (mean = −0.0181) and high-anhedonia subjects essentially balanced (mean = −0.0005), a significant shift in the top-down direction (β = +0.0161, 95% CI [+0.0091, +0.0231], p < 0.001; d = +0.71; R² = 0.133) that was robust across sensitivity models and on the unnormalized difference (β = +0.122, p < 0.001).

At finer grain, a theory-driven set of six directed PFC→striatum edges, selected to connect established reward nodes via functionally validated pathways^3,32^, provided an exploratory localization of the effect. The dlPFC→NAc edge was the most significant edge and the only one to survive FDR correction in all three models (β = +0.0110; full q = 0.005; patients q = 0.026; patients + diagnosis q = 0.034), consistent with a shift in the top-down direction along this pathway, although its split-half reliability was modest (see below). Two further edges survived correction in the full model but were marginal under the stricter models: vmPFC→NAc (q = 0.027) and ACC→anterior putamen (q = 0.027). The remaining edges did not survive correction (Supplementary Table 2).

### The top-down shift reflects a loss of ascending drive, not stronger descending drive

The shift in the top-down direction at both boundaries could arise in two ways: the cortex driving the subcortex harder (stronger descending drive), or the subcortex driving the cortex less (diminished ascending drive). Decomposing each asymmetry into its two directed arms (**Figure 1f**) resolved this in favor of the latter. At the cortico-subcortical boundary, both arms were weaker in high anhedonia, an overall reduction in inter-compartmental coupling: descending cortex→subcortex (β = −2.51, p = 0.012; **Figure 5b**) and ascending subcortex→cortex (β = −3.45, p = 0.001; **Figure 5c**). The ascending arm, however, fell substantially more, and it is this differential that tips the net balance toward top-down. The frontostriatal circuit showed the same pattern: the ascending striatum→PFC arm fell significantly (β = −0.42, p = 0.014; **Figure 5f**), while the descending PFC→striatum arm declined only weakly and non-significantly (β = −0.29, p = 0.077; **Figure 5e**). High anhedonia is therefore characterized not by stronger cortical output but by a disproportionate loss of ascending drive, so that cortical influence is no longer outweighed.

### Flow asymmetry scales with anhedonia severity

Finally, we asked whether the directed-flow measures track anhedonia severity at the individual level, correlating each subject’s normalized flow asymmetry with SHAPS total across the pooled sample (**Figure 6a**). All five associations were positive and survived FDR correction within the primary five-test family: greater top-down dominance accompanied higher SHAPS, the same direction as the categorical group difference. The compartment-level measures were strongest (cortex–subcortex r = +0.27, ρ = +0.36, q < 0.001; PFC–striatum r = +0.24, ρ = +0.35, q = 0.001), and the three single edges were weaker but still survived correction (vmPFC→NAc, dlPFC→NAc, ACC→anterior putamen; all q = 0.028). Spearman ρ equalled or exceeded Pearson r for most measures, indicating a monotonic relationship not driven by outliers.

**Figure 6.**
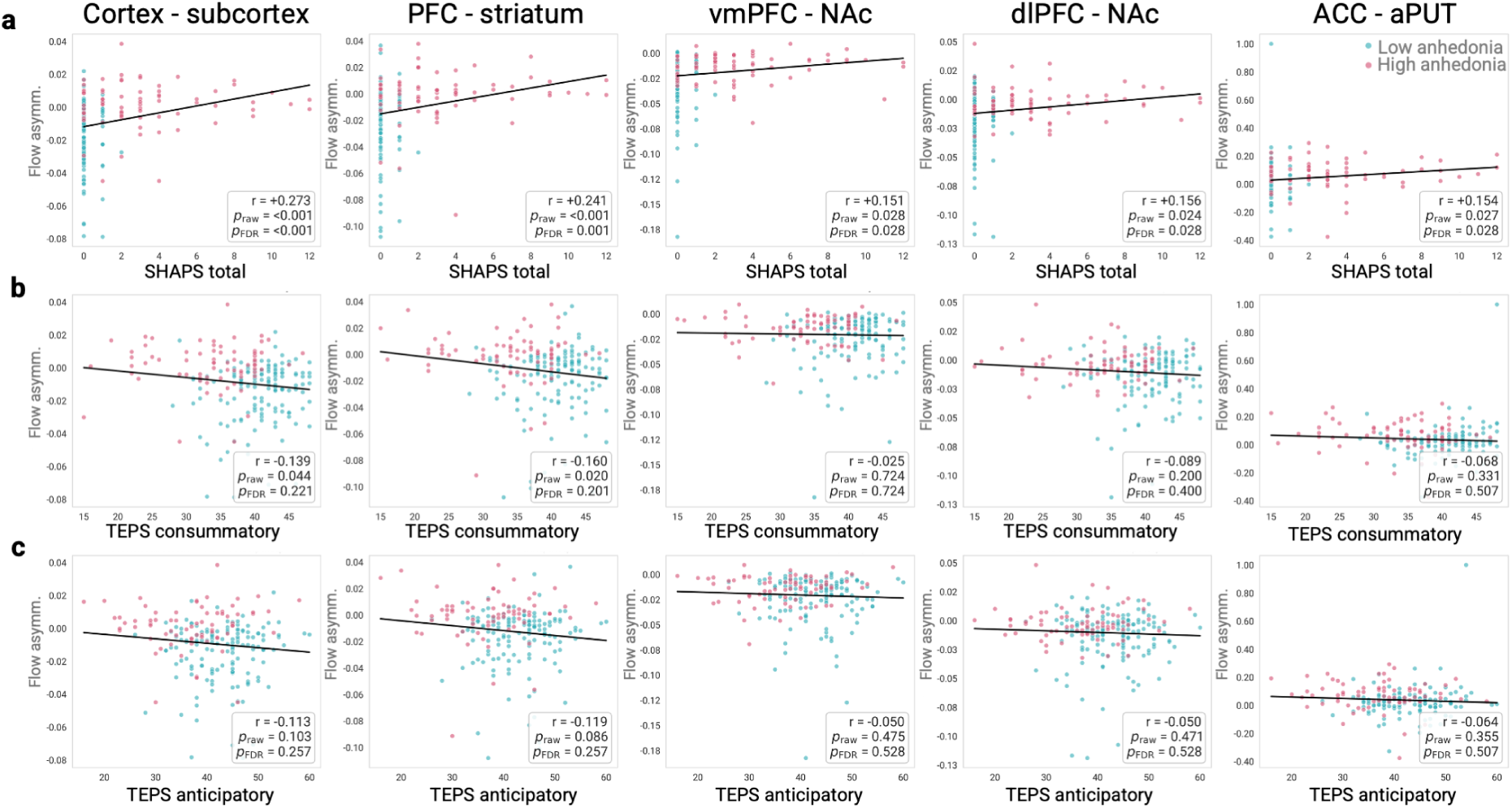
Directed flow asymmetry scales continuously with anhedonia severity. Each subject’s flow-asymmetry index (columns: cortex–subcortex and PFC–striatum compartments, and the vmPFC→NAc, dlPFC→NAc, and ACC→anterior putamen edges) is correlated against three severity measures, with Pearson r and raw/FDR p annotated. Dots are individual participants (n = 210) and the line is the least-squares fit across the pooled sample. aPUT, anterior putamen. **(a)** Against SHAPS total: all five associations are positive and significant (greater top-down dominance with higher SHAPS). **(b)** Against the TEPS consummatory subscale: both compartment measures are negative, as expected because higher TEPS denotes greater hedonic capacity, and are significant before but not after FDR correction, while the single edges show no association. **(c)** Against the TEPS anticipatory subscale: no association survives, localizing the severity relationship to the consummatory facet.

We repeated the analysis with the TEPS, which is scored in the opposite direction (higher TEPS denotes greater hedonic capacity), so that a negative correlation would indicate the same severity relationship as the positive SHAPS correlation. The compartment-level measures correlated negatively with TEPS total in the expected direction (cortex–subcortex r = −0.14, ρ = −0.17; PFC–striatum r = −0.16, ρ = −0.20), matching the categorical group contrast, but neither survived FDR correction within the secondary family (both q = 0.091). Decomposing the TEPS into its subscales, the consummatory rather than the anticipatory facet showed the numerically stronger association for both compartment measures (consummatory cortex–subcortex r = −0.14, PFC–striatum r = −0.16; anticipatory both |r| ≤ 0.12), but none survived correction within the specificity family (all q ≥ 0.20; **Figure 6b-c**), and none of the three frontostriatal edges correlated with TEPS in any form (all q ≥ 0.34; Supplementary Table 3). The TEPS associations are therefore best read as a same-direction but sub-threshold trend: the dimensional severity relationship is carried by SHAPS, with TEPS convergent in sign but not surviving multiple-comparison correction.

### The signature is only weakly reflected in undirected functional connectivity

To test whether these effects depend on directionality, the key group analyses were repeated using static functional connectivity (the Fisher-z–transformed Pearson correlation between parcellated regional time series) in place of the GEC-derived measures, under the same covariate-adjusted design, mapping each directed measure to its closest undirected analogue (Methods). No functional connectivity comparison reached significance in any family, and the directed signature was at most partially echoed. The directed analysis had produced two distinct findings at the cortico-subcortical boundary, and functional connectivity behaved differently toward each.

The first was the hierarchical flattening, cortex shifting upstream relative to subcortex, which the trophic gap captured as the study’s single strongest effect. Its undirected analogue is the difference between cortical and subcortical mean connectivity strength, and its group difference was essentially zero (β = +0.0006, p = 0.95): cortical and subcortical strength declined together, leaving their difference unchanged, whereas the directed gap fell to about zero. The second finding was a reduction in the overall magnitude of coupling between the two compartments (in the directed data, both the ascending and descending arms weakened). Its undirected analogue is the mean strength of the connections crossing the boundary, and here functional connectivity trended the same way, though sub-significantly: cross-boundary coupling was lower in high anhedonia at the cortico-subcortical boundary (β = −0.0165, 95% CI [−0.0406, +0.0076], p = 0.18) and, more weakly, at the frontostriatal boundary (β = −0.0077, p = 0.60). The reward circuit told the same story: no individual reward ROI, the PFC–striatum strength gap, or any of the six frontostriatal edges reached significance, and the strongest directed edge, dlPFC↔NAc, was flat in functional connectivity (β = −0.0002, p = 0.99).

The anhedonia-related signal is thus expressed far more sensitively in directed hierarchy than in undirected connectivity. Because functional connectivity is symmetric, it registers changes in coupling magnitude but not in direction: it retained a faint, same-sign trace of the overall drop in cross-boundary coupling, but none of the hierarchical reorganization, the cortex–subcortex hierarchical flattening and its flow-asymmetry counterpart, that carried the significant, reproducible effects (Supplementary Discussion; Supplementary Table 4; Supplementary Fig. 2).

### Compartment-level effects replicate across independent halves of the data

To assess stability, we refitted each participant’s GEC separately on two halves of their data (run 1 and run 2, each comprising both phase-encoding directions) and recomputed every measure (Methods). The compartment-level measures were moderately to highly reliable across halves (Spearman–Brown-corrected split-half reliability: cortex–subcortex gap 0.80, cortex–subcortex flow index 0.83, PFC–striatum gap 0.76, PFC–striatum flow index 0.82), and the group effect on each replicated independently within both halves (all p ≤ 0.004; Supplementary Table 5). The null result for global trophic coherence also replicated. The single-edge measures were less reliable (Spearman–Brown 0.39–0.68), and the dlPFC→NAc effect was significant in one half (p = 0.001) but not the other (p = 0.09). We therefore treat the edge-level results as exploratory localization of the compartment-level effect.

## Discussion

We asked whether the directed, hierarchical organization of resting-state brain dynamics differs with transdiagnostic anhedonia, and found a specific signature. Global trophic coherence did not differ between low- and high-anhedonia groups. Instead, the hierarchy was redistributed: in high anhedonia the cortex shifted upstream relative to the subcortex, eliminating the bottom-up ordering seen in low anhedonia. The cortex–subcortex hierarchical gap, positive in low anhedonia (cortex downstream of subcortex), fell to about zero in high anhedonia. This flattening of the cortico-subcortical hierarchy was strongest at the level of whole compartments, was mirrored within the reward circuit (a PFC–striatum gap that likewise fell to about zero and, at the edge level, a top-down shift along dlPFC→NAc), scaled with anhedonia severity across individuals, and was only weakly and non-significantly reflected in undirected functional connectivity. Taken together, these results identify a directed-hierarchy reorganization of cortico-subcortical and frontostriatal circuits as a candidate transdiagnostic substrate of anhedonia.

Within the reward circuit, the frontostriatal pattern maps onto the spatial pattern predicted by the dominant neurobiological model of anhedonia. A large body of reward-task and resting-state work converges on striatal hypofunction alongside over-recruitment of medial and dorsolateral prefrontal cortex, an imbalance those studies interpret as compensatory top-down control over a blunted reward response^3,10,11,15,33–35^. Our findings provide what is, to our knowledge, the first directionally-resolved, whole-brain view of that frontostriatal imbalance at rest: when each region’s position is read out from a generative, directed model of brain dynamics, the prefrontal components of the reward circuit – the OFC, a principal reward–hedonic hub^36,37^, together with mPFC/vmPFC, ACC, and dlPFC – occupy a more upstream position relative to the striatum than in low anhedonia. The strongest edge was dlPFC→NAc, the corticostriatal pathway by which the prefrontal cognitive-control system most directly influences the ventral striatum^38,39^ and thus the edge most naturally identified with the top-down control posited by over-recruitment accounts^3^. That the prefrontal effects grew statistically stronger as healthy controls and diagnostic category were partialled out indicates that they track anhedonia severity rather than clinical status or diagnosis, consistent with anhedonia’s status as a dimensional, transdiagnostic symptom. That the strongest edge was dlPFC→NAc, an established frontostriatal pathway with demonstrated functional relevance^32^, is consistent with the effect being expressed along canonical reward-circuit connections, although edge-level measures were less reliable across halves of the data than compartment-level ones, so this localization should be treated as exploratory. The hierarchical gap and the flow-asymmetry index are both computed from the same GEC weights. For cortex versus subcortex, which together make up the whole brain, an edge-weighted version of the gap is mathematically equivalent to the flow index, and the unweighted gap we report closely tracks it (r = −0.94; Methods). For smaller compartments such as PFC and striatum, no such equivalence holds, so their agreement there shows that the shift in hierarchical position is matched by a shift in the direct flow balance between the two. However, reproducing this pattern is not the same as confirming the mechanism that the over-recruitment account proposes for it.

Our result matches the direction of change the account predicts, a shift in prefrontal–striatal balance toward the prefrontal side, but, as we show next, the directed decomposition reveals that this shift arises for a different reason.

Decomposing the boundary flow into its ascending and descending arms revealed that high anhedonia is not characterized by cortex driving subcortex harder in absolute terms; rather, both directed flows were weaker, and the ascending (subcortex→cortex, striatum→PFC) arm declined disproportionately more than the descending arm. The net shift away from bottom-up dominance is therefore produced by a disproportionate reduction in ascending drive, not by an absolute increase in prefrontal output. In low anhedonia, ascending drive from subcortex and striatum outweighs descending prefrontal influence; in high anhedonia, that predominance is lost, so prefrontal influence is no longer outweighed even though it has not grown. This reframes the “prefrontal over-recruitment” seen in activation studies (**Figure 7**): at the level of directed network flow, what appears as excess top-down control may instead reflect the loss of the normal bottom-up predominance of reward and salience signaling from subcortex to cortex^3,40–43^. The distinction matters mechanistically and therapeutically, because it locates the deficit in the ascending drive of the circuit rather than in prefrontal hyperactivity per se.

**Figure 7.**
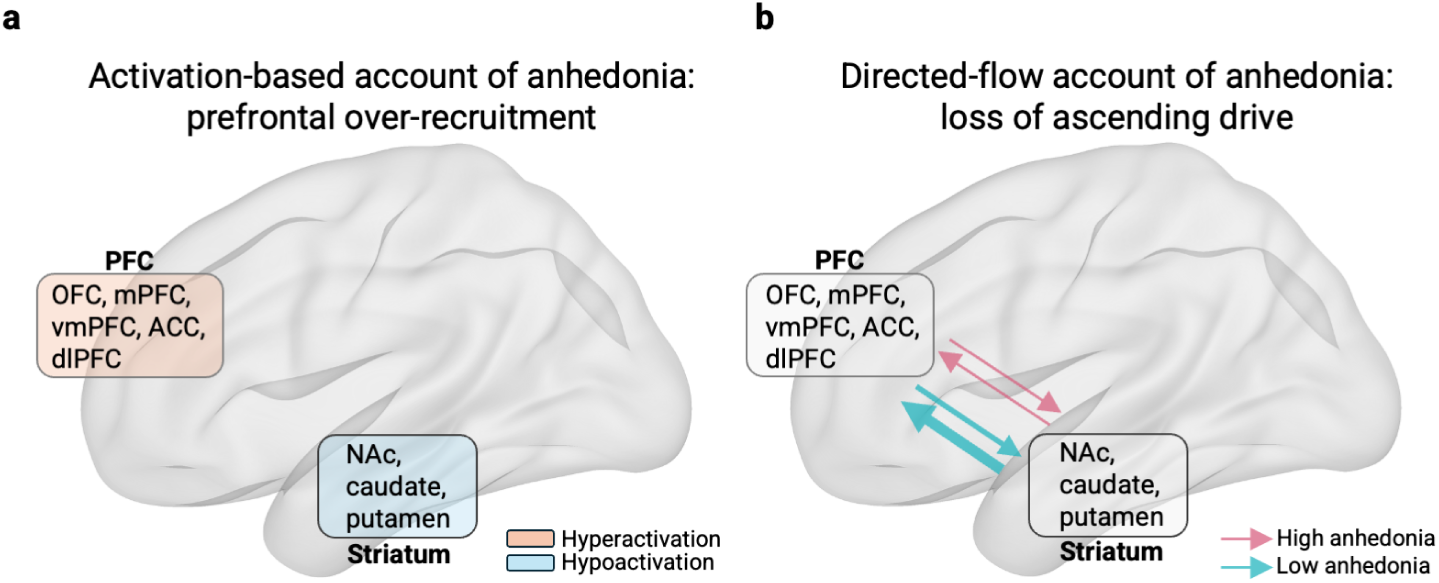
Contrasting accounts of frontostriatal imbalance in anhedonia. **(a)** Previous activation-based research has associated anhedonia with an imbalance within the reward circuit, in which activation is relatively increased in prefrontal regions and decreased in the striatum, a pattern interpreted as compensatory prefrontal over-recruitment^33^. **(b)** The directed-coupling picture from the present study, over the same circuit: GEC weight between the prefrontal (OFC, mPFC/vmPFC, ACC, dlPFC) and striatal (NAc, caudate, putamen) compartments in the low- and high-anhedonia groups, with arrow thickness proportional to the group-mean GEC weight. Both the descending (PFC→striatum) and ascending (striatum→PFC) arms are weaker in high anhedonia, but because the ascending arm declines disproportionately, the net balance shifts from bottom-up to balanced. At the level of directed coupling there is therefore no evidence of increased prefrontal drive: what appears in **(a)** as excess top-down control may instead reflect diminished ascending drive.

Beyond the reward circuit, all seven Yeo cortical networks were lower relative to each participant’s most upstream region (largest in somatomotor and default-mode cortex, networks with well-documented functional-connectivity abnormalities in depression^44^), although no single network survived multiple-comparison correction. Because these network-level differences depend on the choice of reference (Methods), we base our interpretation on the cortex–subcortex gap, which does not: a change in the cortex’s position relative to the subcortex within the directed hierarchy, consistent with evidence that the brain’s directed hierarchy reconfigures across brain states^25^ and with proposals linking disruption of the pleasure system to altered whole-brain dynamics and metastability^45^.

These results reinforce a dimensional, transdiagnostic reading of anhedonia. High anhedonia was present across virtually every diagnostic category yet enriched in mood and psychotic disorders, the flow-asymmetry measures scaled continuously with severity across two oppositely-scored instruments (SHAPS and TEPS), and defining anhedonia from a multi-scale factor solution rather than a single cut-off captured variance that any one instrument would have missed^4^. More broadly, this dimensional, transdiagnostic reading is consistent with the RDoC framework’s emphasis on neurobiologically grounded dimensions that cut across DSM boundaries^8^ and with transdiagnostic neuroimaging work linking anhedonia to frontostriatal and vmPFC–ventral-striatal dysfunction independent of diagnosis^9–11^. The dimensional severity relationship was carried by SHAPS, which correlated with all five flow measures after correction; the TEPS reproduced the same direction but did not survive multiple-comparison correction. The numerically stronger TEPS associations fell on the consummatory rather than the anticipatory subscale – a distinction grounded in the separation of anticipatory, motivational “wanting” from consummatory, hedonic “liking” components of reward^4,12,13,17,29,46–48^. This consummatory bias aligned in sign with the predominantly consummatory SHAPS, suggesting, without establishing, that the cortico-subcortical and frontostriatal flow rebalancing may track consummatory more than anticipatory anhedonia. Wanting and liking are, however, only two phases of a repeating pleasure cycle whose third phase, satiety, signals when enough has been obtained so that one pursuit can yield to another; selective satiety is the operation that turns hedonic processing into a scheduler of competing needs rather than a single reward value^12,49,50^. Neither the SHAPS nor the TEPS assesses satiety, so our data cannot determine which phase of the cycle the directed-flow changes track. The same framework also allows anhedonia to arise not only from disturbed wanting or liking but from altered large-scale orchestration that subsequently deforms affective scheduling^50^, a possibility consistent with the directed-hierarchy reorganization observed here.

Several limitations warrant discussion. First, the design is cross-sectional and correlational; unlike before/after intervention designs that can attribute hierarchical reconfiguration to a manipulation^20,51^, we cannot infer whether the directed-flow signature is a cause, consequence, or correlate of anhedonia. Second, although age and sex were balanced and patient/healthy-control status was modeled as a covariate, residual confounding from clinical status cannot be fully excluded, because anhedonia and illness are strongly correlated in this sample and binary patient/control status is only a coarse proxy for illness burden; medication and comorbid secondary diagnoses were likewise not modeled. That said, the principal gap and flow effects strengthened rather than weakened in the patients-only and diagnosis-adjusted sensitivity analyses, arguing against their being a mere by-product of clinical status. Third, the parcellation we used did not resolve every region needed for the reward-circuit analyses, so some canonical nodes could only be approximated. For example, the subcortical atlas contains no ventral tegmental area (VTA)/midbrain parcel (the dopaminergic origin of the mesocorticolimbic circuit) and does not resolve the subgenual (BA25) cingulate, leaving the most ventral reward nodes incompletely represented. Dedicated probabilistic atlases of these nuclei already exist^52,53^, and future work using higher-resolution or more targeted parcellations could capture the full set of nodes and edges the canonical circuit requires.

In sum, by resolving the direction of information flow, this approach reframes prefrontal “over-recruitment” in anhedonia as a relative dominance arising from diminished ascending drive, and identifies cortico-subcortical and frontostriatal flow asymmetry, which tracks anhedonia severity, as a promising target for future modeling. Natural next steps include testing whether neurotransmitter-informed whole-brain modeling can identify the receptor systems whose perturbation most effectively “rescues” the reorganized hierarchy in silico, and whether the baseline signature predicts treatment response in longitudinal designs.

## Methods

### Participants and dataset

Data were drawn from the TCP, an open dataset of 241 adults recruited across Yale University and McLean Hospital^26^, comprising healthy controls and individuals meeting diagnostic criteria across affective, anxiety, and psychotic disorders. Participants completed a clinician interview and a broad behavioral and cognitive battery, and were scanned on harmonized 3T Siemens Prisma systems (dataset and analysis-pipeline overview, **Figure 1**). Analyses used the subset of participants with a derived anhedonia cluster assignment and complete resting-state imaging and demographic data; sample sizes for each model are reported in Results. The TCP has been analyzed previously with undirected network-dynamics methods in an overlapping sample (n = 219)^27^, which used resting-state and task fMRI to classify multi-dimensional symptom profiles; neither directed connectivity nor anhedonia was examined there. All data were fully de-identified prior to release. Original TCP data collection was approved by the Institutional Review Boards of Yale University and McLean Hospital, and all participants provided written informed consent^26^. The present secondary analysis of de-identified data was carried out under an approved NIMH Data Archive (NDA) Data Use Certification (data access request DAR-25575).

### Anhedonia measures, factor analysis, and clustering

Anhedonia was operationalized transdiagnostically from three self-report scales in the TCP battery: the 14-item SHAPS^28^, the 18-item TEPS^29^ (anticipatory and consummatory subscales), and a five-item anhedonia composite from the DASS^30^. Missing item responses were handled by multiple imputation by chained equations (MICE) in R after excluding participants with more than 60% missingness within any scale. Because the items are ordinal or binary, all factor-analytic computations were performed on a polychoric correlation matrix. Suitability was confirmed with Bartlett’s test of sphericity (χ² = 3985.68, p < 0.001) and the Kaiser–Meyer–Olkin measure (KMO = 0.852). The number of factors was selected by Horn’s parallel analysis (observed eigenvalues versus the 95th percentile of eigenvalues from 200 random datasets of matched dimensions), which supported four factors. Factors were extracted by principal axis factoring with varimax rotation, and per-subject factor scores were computed by the regression method.

Participants were clustered into two anhedonia subgroups by fitting a Gaussian mixture model to the four standardized factor scores, yielding a low-anhedonia group (n = 145) and a high-anhedonia group (n = 74). The solution was validated with Welch’s t-tests on the three scale totals (FDR-corrected across the three comparisons) and characterized by the distribution of cluster membership across primary diagnoses. These subgroups are the grouping variable (reference = low anhedonia) for all neuroimaging analyses.

### fMRI acquisition, preprocessing, and parcellation

Resting-state fMRI was acquired on 3T Siemens Prisma systems (2-mm isotropic, TR = 800 ms, multiband factor 8), with four runs per participant (run 1/2 × AP/PA phase encoding). We used the HCP minimally-preprocessed, ICA-FIX–denoised CIFTI files (*_Atlas_MSMAll_hp2000_clean.dtseries.nii, without global signal regression). The four runs were parcellated and the time series concatenated in the order run1-AP, run1-PA, run2-AP, run2-PA. Parcellation combined the Schaefer-200 (7-network) cortical atlas^54^ with the Tian Scale-II 32-region subcortical atlas^55^, giving 232 regions (indices 0–31 subcortex, 32–231 cortex), the convention used throughout.

### Generative effective connectivity

For each subject we estimated a directed, weighted GEC matrix (232 × 232) within the whole-brain Hopf modeling framework^19–21^ (**Figure 1d**). Each region *i* was modeled as a Stuart–Landau (Hopf) oscillator – the normal form of a supercritical Hopf bifurcation – with complex state *z_i_* = *x_i_* + *i y_i_*, coupled to the other regions through the connectivity matrix *C*:

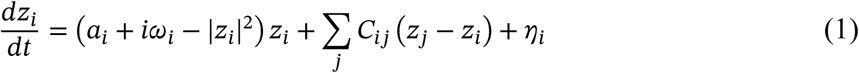

where *η_i_* is additive uncorrelated Gaussian noise with standard deviation *σ* = 0.01 (homogeneous across regions, so the noise covariance is *Q* = *σ*^2^*I*), and the simulated BOLD signal is the real part *x_i_* = Real(*z_i_*). The bifurcation parameter *a_i_* sets the local regime: for *a_i_* < 0 the region settles into a stable, noise-driven fixed point, and for *a_i_* > 0 it engages a self-sustained oscillation of frequency *ω_i_*/2*π*. All regions were held just below the bifurcation at *a_i_* = −0.02, the near-critical working point at which the model best reproduces empirical resting-state dynamics. The intrinsic frequency *ω_i_* = 2*πf_i_* was fixed per region from the data, with *f_i_* the peak frequency of that region’s band-pass-filtered BOLD signal.

Because the dynamics sit near the stable fixed point *z* = 0, the system can be linearized, giving an analytic solution for the model covariance and avoiding stochastic simulation. In terms of the

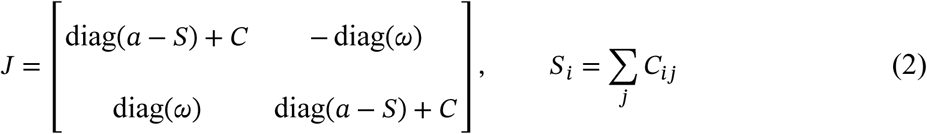

the stationary covariance *K* (2*N* × 2*N*) solves the continuous Lyapunov equation

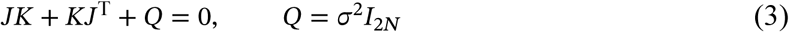

and the model zero-lag functional connectivity FC^model^ is the correlation matrix of the upper-left *N* × *N* block of *K* (the real, BOLD-related part). The model *τ*-shifted covariance follows in closed form as KS^model^(*τ*) = exp(*τJ*) ⋅ *K*, evaluated at a lag *τ* = 3 TR = 2.4 s and normalized as below to give FS^model^(*τ*).

The anatomical scaffold for the model was the probabilistic tractography-based normative connectome distributed with the Lead-DBS toolbox^56^ (https://www.lead-dbs.org/). That connectome was derived from diffusion-weighted and T2-weighted MRI in 32 healthy participants (mean age 31.5 ± 8.6 years, 14 female) from the Human Connectome Project^57^, whose diffusion data were acquired over 89 minutes on a scanner with stronger gradients than conventional systems (acquisition details available through the Image & Data Archive, https://ida.loni.usc.edu/). A generalized q-sampling imaging algorithm was applied to the diffusion data in DSI Studio (http://dsi-studio.labsolver.org). Images were co-registered to the b0 image of the diffusion data within a white-matter mask segmented from the T2-weighted images using SPM12, and the 200,000 most probable fibres per participant were sampled inside that mask. Fibres were then warped to MNI space using a nonlinear deformation field derived from the T2-weighted images^58^, and the individual tractograms were aggregated into a single normative tractogram representative of a healthy young-adult population^56^. The same normative connectome has been used as the coupling structure in whole-brain oscillator models of this kind^59^.

From this normative tractogram a single group-average weight matrix was computed over the combined Schaefer-200 (7-network) cortical and Tian Scale-II subcortical parcellation used throughout this study (232 regions)^54,55^ in MNI152NLin2009cAsym space, with each entry *C*(*n*, *p*) given by the number of streamlines running between the voxels belonging to regions *n* and *p*. Because the linearized model used here treats interactions as instantaneous, only this weight matrix was carried forward; the corresponding matrix of mean tract lengths, which is required only when conduction delays are modelled explicitly, was not used.

The oscillators were coupled through a matrix *C* initialized from this group-average structural connectome, rescaled so its maximum weight was 0.2. *C* was refined by a heuristic pseudo-gradient procedure that updates only connections present in the anatomical scaffold within each hemisphere, with the exception that homologous inter-hemispheric connections between the same region in either hemisphere were also updated, because tractography underestimates these.

The empirical targets were computed from each subject’s concatenated parcellated time series after demeaning, linear detrending, and zero-phase band-pass filtering (second-order Butterworth, 0.01 Hz to 0.99 × the Nyquist frequency, ≈0.62 Hz at TR = 0.8 s), trimming the first and last 10 volumes: the empirical functional connectivity FC^emp^ as the Pearson correlation matrix and the *τ* time-shifted covariance at *τ* = 3 TR. The fit drove the model to reproduce both simultaneously.

The time-shifted covariance was normalized by dividing each pair (*i*, *j*) by the geometric mean of its zero-lag variances,

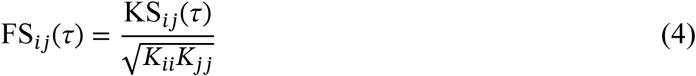

a normalization that breaks the symmetry of the couplings and allows the procedure to recover directionality. Unlike the arrow-of-time formulation of Deco et al.^20^, this approach does not fit a forward-versus-time-reversed contrast; it matches the forward time-shifted covariance directly. At each iteration the coupling was updated by

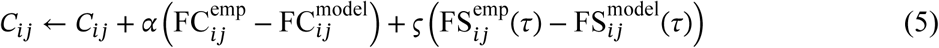

with the model matrices FC^model^ and FS^model^(*τ*) obtained in closed form from the linearized system (the Lyapunov solution *K* and KS^model^(*τ*) = exp(*τJ*) ⋅ *K* above), and learning rates *α* = 4 × 10^−4^ for the zero-lag term and *ς* = 1 × 10^−4^ for the time-shifted term. After each update, negative weights were set to zero, *C* was rescaled to a maximum weight of 0.2, and only anatomically supported entries were modified. Iteration continued until the summed FC and FS(*τ*) error ceased to improve (relative change < 0.1% over 100 iterations; maximum 5000 iterations). Fitting proceeded in two stages: a group-level *C*_eff_ was first estimated from the group-average empirical targets (seeded from the structural connectome), and each subject’s *C*_eff_ was then estimated by the same procedure seeded from that group solution; low- and high-anhedonia subjects were fit with identical settings. Fit quality was quantified as the Pearson correlation between empirical and model matrices over their lower triangles. The resulting *C*_eff_ is asymmetric, with the convention *C*_eff_[*i*, *j*] = directed influence from region *j* to region *i* (row = target, column = source). GEC matrices were estimated in MATLAB and loaded as precomputed files for the analyses reported here.

### Trophic levels and trophic coherence

Hierarchical organization was read out from each subject’s *C*_eff_ using trophic levels in the basal-node-free, reversal-symmetric formulation of MacKay et al.^24^, as adopted by Deco et al.^20^. Treating *C*_eff_ as a directed weighted graph with 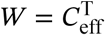, so that *W_ij_* is the weight from region *i* to region *j*, we defined each region’s in-weight and out-weight as the column and row sums of *W*, the total weight *u* = in + out, and the imbalance *v* = in − out. Trophic levels *h* solve the symmetric graph-Laplacian system

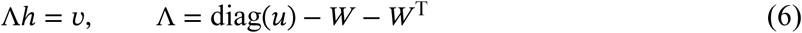

with the gauge fixed by anchoring one node; levels were then shifted so the minimum is 0. Unlike the classical ecological definition, this formulation requires no basal node and treats forward and reverse flow symmetrically, so it can be computed on a fully recurrent connectome. A region’s trophic level reflects its position in the global directed hierarchy and depends on the entire network, including indirect multi-hop pathways: higher levels denote downstream positions, where incoming GEC weight exceeds outgoing, lower levels upstream positions, where the balance reverses. Because the gauge is fixed by anchoring a single node, levels carry no absolute meaning and are interpreted only relative to one another, so group differences index a change in relative position rather than in absolute coupling magnitude. Because the region with the minimum level differs across participants, a group difference in a single region’s, network’s or compartment’s mean level describes its position relative to each participant’s most upstream region and depends on this choice of reference. Our primary hierarchical outcomes are therefore the within-participant differences between compartments (the cortex–subcortex and PFC–striatum gaps), which are independent of the reference. We accordingly describe regions as upstream or downstream rather than in the absolute source/sink terms common in ecology research. The same solution yields the scalar trophic coherence (directedness) *H* = 1 − *F*_0_, where *F*_0_ is the edge-weight-normalized mean squared deviation of level differences from the ideal value of 1,

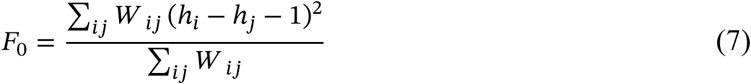

Trophic levels and coherence were also computed on the cortical-only (200 × 200) and subcortical-only (32 × 32) submatrices for compartment-specific analyses.

### Compartment, network, and reward-circuit summaries

From each subject’s regional trophic-level vector we derived several summaries. At the compartment level, we averaged trophic levels within cortex and within subcortex and computed their difference (the cortex–subcortex hierarchical gap). At the network level, we averaged within each of the seven Yeo cortical networks plus a subcortical “network” (FDR across the eight). For the reward circuit, we grouped regions into seven ROIs defined from the Schaefer/Tian labels: three striatal (NAc, caudate, putamen) and four cortical (OFC, mPFC/vmPFC, ACC, dlPFC), and additionally collapsed these into a prefrontal compartment (OFC + mPFC/vmPFC + ACC + dlPFC) and a striatal compartment (NAc + caudate + putamen) with their difference (the PFC–striatum gap). Three atlas caveats apply to the reward analyses: the Tian-S2 atlas contains no VTA/midbrain parcel (the dopaminergic origin of the mesocorticolimbic circuit), does not resolve subgenual (BA25) cingulate (perigenual/ventromedial signal falls within the mPFC/vmPFC ROI; the ACC ROI reflects dorsal/mid-cingulate), and merges medial and lateral OFC.

### Cortico-subcortical and frontostriatal flow asymmetry

In addition to global trophic positioning, we computed a direct measure of net directed flow across two boundaries. For the cortico-subcortical boundary we summed the *C*eff weights for cortex→subcortex (*A*) and subcortex→cortex (*B*) and formed both the raw difference (top-down dominance = *A* − *B*; positive = top-down) and a normalized index (*A* − *B*)/(*A* + *B*), bounded in [−1, +1]. The normalized index removes the influence of overall coupling magnitude, comparing the balance rather than the size of flow; group tests used the normalized index and were confirmed on the unnormalized difference. To interpret any group difference, the asymmetry was decomposed into its two arms (descending and ascending tested separately), revealing whether the arms moved in the same or opposite directions. The same logic was applied to the frontostriatal sub-network (PFC↔striatum). At finer grain we tested a theory-driven set of six directed PFC→striatum edges (vmPFC–NAc, OFC–NAc, ACC–NAc, dlPFC–NAc, vmPFC–anterior putamen, ACC–anterior putamen), each retained only if the striatal node is a reward node per Pizzagalli3 and the connection is a functionally validated pathway per Morris et al.32; the anterior putamen was used for the dorsal-striatal edges. FDR correction was applied across the six edges within each model, and the two-arm decomposition was run for each FDR-significant edge.

The two readouts are mathematically related. Trophic levels are solved from the balance of incoming and outgoing weight at every region, so a compartment’s mean level is tied to its net flow with the rest of the network. When two compartments together make up the whole network, as cortex and subcortex do, the relationship is exact: summing equation 6 over all cortical regions cancels every within-cortex term, leaving ∑_*i∈ctx*, *j∈sub*_(*W _ij_* + *W _ji_*)(*h_i_* − *h_j_*) = *B* − *A*, so the cortex–subcortex level difference weighted by the cross-boundary edges equals minus the normalized flow index. The unweighted gap we report is not identical to this quantity but closely tracked the index across participants (r = −0.94). No such identity holds for PFC and striatum, which do not partition the network, so agreement there is not guaranteed by construction. We therefore use the flow index not as independent confirmation of the gap but for what it adds: its two arms can be tested separately, attributing a shift in balance to a change in ascending or descending drive.

### Statistical modeling and covariates

Group balance was checked before the imaging analyses: age (continuous) with a Wilcoxon rank-sum test, reported as the normal-approximation Z; sex and patient/healthy-control status (categorical) with χ² tests of independence on 2×2 tables using Yates’ continuity correction. Participants missing age or reporting a sex category other than male/female were excluded from the corresponding test only and retained in all imaging analyses.

Because this confound check showed the groups were imbalanced on clinical status, every neuroimaging metric was tested with OLS regression rather than a raw group contrast:

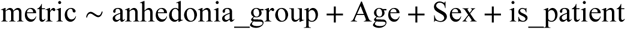

with reference levels low-anhedonia, female, and healthy control. The effect of interest is the coefficient for the high-versus-low anhedonia contrast after partialling out the covariates. Age and sex were included as standard prognostic covariates that reduce residual outcome variance, while clinical status was included primarily to address its confounding with anhedonia group. Each metric was additionally probed with two sensitivity models: (2) patients only (healthy controls excluded, is_patient dropped), and (3) patients only plus a broad three-level diagnostic covariate (Mood / Anxiety-Trauma / Other). Specific diagnoses were not entered individually, both for cell-count reasons and because diagnoses such as MDD are defined partly by anhedonia. Where a family of related metrics was tested (regions, networks, ROIs, edges), p-values were Benjamini–Hochberg FDR-corrected across the family. To test whether each group’s gap and flow index differed from zero, we refitted the same model with one coefficient per group and covariates centred at their sample means, so that each group coefficient is that group’s covariate-adjusted mean. As a standardized effect size for the compartment-level measures, we report d, the covariate-adjusted group coefficient divided by the pooled within-group standard deviation of the outcome.

### Dimensional severity associations

To test whether the directed-flow measures track severity dimensionally rather than only between clusters, each normalized flow-asymmetry index (the cortico-subcortical and frontostriatal compartment indices and the three frontostriatal edges that survived the group analysis) was correlated with continuous symptom totals across the pooled sample (both groups; N = 210), reporting both Pearson r and Spearman ρ. SHAPS total and TEPS total were computed by summing the imputed item scores; the TEPS was additionally decomposed into its anticipatory (items 1, 4, 6, 8, 10, 11, 13, 15, 16, 18) and consummatory (items 2, 3, 5, 7, 9, 12, 14, 17) subscales to test which facet of anhedonia the flow measures track. The two instruments are scored in opposite directions (higher SHAPS denotes greater anhedonia, higher TEPS greater hedonic capacity), so a positive SHAPS correlation and a negative TEPS correlation each indicate that more top-down-dominant flow accompanies greater anhedonia. Because these correlations span five brain measures and up to four symptom scores, p-values were Benjamini–Hochberg FDR-corrected within hypothesis-defined families rather than pooled: a primary family of each measure’s correlation with SHAPS total (the a priori primary hedonic-capacity scale; five tests), a secondary family of the same five measures against TEPS total (convergent validity), and a specificity family of the five measures against the two TEPS subscales (ten tests). Pearson correlations were corrected as the primary read-out, with Spearman ρ reported as a sensitivity check.

### Functional-connectivity control analysis

To assess whether the findings are recoverable from undirected connectivity, the key group analyses were repeated using static functional connectivity, defined as the Fisher-z–transformed Pearson correlation between the parcellated regional time series, in place of the GEC-derived measures and under the identical covariate-corrected OLS design and the same three-model and FDR conventions. Because functional connectivity is symmetric and edge-based rather than directed and nodal, each GEC measure was mapped to its closest undirected analogue: a region’s or ROI’s trophic level (a nodal, directional quantity) was matched by its nodal functional connectivity strength (mean functional connectivity to all other regions), and each directional flow asymmetry across a boundary (cortex↔subcortex, PFC↔striatum) or along a directed edge was matched by the corresponding mean undirected functional connectivity across that boundary or ROI pair (a single value, with no ascending/descending arms to decompose). The same six families tested for trophic levels and flow asymmetry were re-run on the functional connectivity matrices. This comparison was intended as a matched analogue of the directed analyses rather than as an optimized test of connectivity-based prediction in this cohort (cf. Cocuzza et al.27); feature definition, covariates, model structure, and FDR conventions were therefore held identical across the directed and undirected analyses, and no dimensionality-reduction or multivariate-learning step was applied to either. No functional connectivity metric reached significance in the covariate-adjusted model and none survived FDR correction in any family, though cross-boundary coupling trended weaker in high anhedonia in the same direction as the directed reduction; full results are reported in the Supplementary Discussion (Supplementary Table 4; Supplementary Fig. 2).

### Split-half reliability

To estimate the stability of the measures, we split each participant’s four runs into two halves of equal length (half 1: run1-AP + run1-PA; half 2: run2-AP + run2-PA) and refitted the GEC separately on each half using the same procedure and settings as the main analysis; as there, each fit was initialised from the group-level GEC and regional intrinsic frequencies estimated from all four runs. All measures were recomputed from each half. Reliability was quantified as the observed Pearson correlation between halves, its Spearman–Brown correction (2r/(1 + r)), which estimates the reliability of the full four-run measure, and the absolute-agreement intraclass correlation ICC(A,1). The primary group model was re-run separately within each half.

### Use of large language models

A large language model (Claude, Anthropic) was used for assistance with manuscript writing and editing, with code development for the analyses, and with checking the manuscript against the journal’s submission guidelines. All text and code produced with this assistance were reviewed, verified and revised by the authors, who take full responsibility for the content of this manuscript.

### Data availability

Data are from the Transdiagnostic Connectome Project26 (NDA Study #3397; https://doi.org/10.15154/w59r-ef41), distributed through the NIMH Data Archive within NDA Collection 3552 (“Functional genomics of the human connectome in psychiatric illness”; https://nda.nih.gov/). The data are available to qualified investigators who obtain an executed NDA Data Use Certification.

### Code availability

All custom code used in this study is available at https://github.com/payman21/tcp-anhedonia-hierarchy and is released under the GNU General Public License v3.0. The repository contains the multiple-imputation scripts for the self-report scales, the anhedonia factor-analysis and clustering notebooks, the fMRI parcellation pipeline, the MATLAB implementation of the Hopf-model generative effective connectivity fitting and the trophic-level decomposition, and the notebooks reproducing all reported statistics and figures.

The repository also includes the group-level effective connectivity matrices and the regional and summary statistics underlying the reported results. Subject-level derived data, including the per-subject effective connectivity matrices, are not redistributed, because the NIMH Data Archive Data Use Certification governing access to these data does not permit redistribution of subject-level derived data. Qualified investigators who obtain an executed Data Use Certification can regenerate them from the source data using the code provided.

## Acknowledgements

Data and/or research tools used in the preparation of this manuscript were obtained from the National Institute of Mental Health (NIMH) Data Archive (NDA). NDA is a collaborative informatics system created by the National Institutes of Health to provide a national resource to support and accelerate research in mental health. Dataset identifier(s): NIMH Data Archive Collection 3552 and NDA Study DOI 10.15154/w59r-ef41. This manuscript reflects the views of the authors and may not reflect the opinions or views of the NIH or of the Submitters submitting original data to NDA.

## Author contributions

P.R. conceived the study, performed the analyses, prepared the figures and wrote the initial manuscript. J.V. contributed to the analysis framework, proposed additional analyses, and contributed to the refinement of the analyses and to the interpretation, and edited the manuscript. M.L.K. contributed to the interpretation and edited the manuscript. G.D. supervised the project, contributed the whole-brain modeling framework and GEC methodology. All authors approved the final manuscript.

## References

1. Guineau, M. G. et al. Anhedonia as a transdiagnostic symptom across psychological disorders: a network approach. Psychol. Med. 53, 3908–3919 (2023).

2. Husain, M. & Roiser, J. P. Neuroscience of apathy and anhedonia: a transdiagnostic approach. Nat. Rev. Neurosci. 19, 470–484 (2018).

3. Pizzagalli, D. A. Toward a better understanding of the mechanisms and pathophysiology of anhedonia: are we ready for translation? Am. J. Psychiatry 179, 458–469 (2022).

4. Rømer Thomsen, K., Whybrow, P. C. & Kringelbach, M. L. Reconceptualizing anhedonia: novel perspectives on balancing the pleasure networks in the human brain. Front. Behav. Neurosci. 9, 49 (2015).

5. Ducasse, D. et al. Anhedonia is associated with suicidal ideation independently of depression: a meta-analysis. Depress. Anxiety 35, 382–392 (2018).

6. Nogo, D. et al. The effect of ketamine on anhedonia: improvements in dimensions of anticipatory, consummatory, and motivation-related reward deficits. Psychopharmacology 239, 2011–2039 (2022).

7. Serretti, A. Anhedonia: current and future treatments. PCN Rep. 4, e70088 (2025).

8. Insel, T. et al. Research Domain Criteria (RDoC): toward a new classification framework for research on mental disorders. Am. J. Psychiatry 167, 748–751 (2010).

9. Geller, W. N., Liu, K. & Warren, S. L. Specificity of anhedonic alterations in resting-state network connectivity and structure: a transdiagnostic approach. Psychiatry Res. Neuroimaging 317, 111349 (2021).

10. Schaub, A.-C. et al. Neural mapping of anhedonia across psychiatric diagnoses: a transdiagnostic neuroimaging analysis. NeuroImage Clin. 32, 102825 (2021).

11. Sharma, A. et al. Common dimensional reward deficits across mood and psychotic disorders: a connectome-wide association study. Am. J. Psychiatry 174, 657–666 (2017).

12. Berridge, K. C. & Kringelbach, M. L. Pleasure systems in the brain. Neuron 86, 646–664 (2015).

13. Der-Avakian, A. & Markou, A. The neurobiology of anhedonia and other reward-related deficits. Trends Neurosci. 35, 68–77 (2012).

14. Haber, S. N. & Knutson, B. The reward circuit: linking primate anatomy and human imaging. Neuropsychopharmacology 35, 4–26 (2010).

15. Keren, H. et al. Reward processing in depression: a conceptual and meta-analytic review across fMRI and EEG studies. Am. J. Psychiatry 175, 1111–1120 (2018).

16. Ng, T. H., Alloy, L. B. & Smith, D. V. Meta-analysis of reward processing in major depressive disorder reveals distinct abnormalities within the reward circuit. Transl. Psychiatry 9, 293 (2019).

17. Treadway, M. T. & Zald, D. H. Reconsidering anhedonia in depression: lessons from translational neuroscience. Neurosci. Biobehav. Rev. 35, 537–555 (2011).

18. Zhang, B. et al. Mapping anhedonia-specific dysfunction in a transdiagnostic approach: an ALE meta-analysis. Brain Imaging Behav. 10, 920–939 (2016).

19. Cabral, J., Kringelbach, M. L. & Deco, G. Functional connectivity dynamically evolves on multiple time-scales over a static structural connectome: models and mechanisms. NeuroImage 160, 84–96 (2017).

20. Deco, G. et al. Different hierarchical reconfigurations in the brain by psilocybin and escitalopram for depression. Nat. Ment. Health 2, 1096–1110 (2024).

21. Kringelbach, M. L. & Deco, G. Brain states and transitions: insights from computational neuroscience. Cell Rep. 32, 108128 (2020).

22. Patow, G., Martin, I., Sanz Perl, Y., Kringelbach, M. L. & Deco, G. Whole-brain modelling: an essential tool for understanding brain dynamics. Nat. Rev. Methods Primers 4, 53 (2024).

23. Gilson, M., Moreno-Bote, R., Ponce-Alvarez, A., Ritter, P. & Deco, G. Estimation of directed effective connectivity from fMRI functional connectivity hints at asymmetries of cortical connectome. PLoS Comput. Biol. 12, e1004762 (2016).

24. MacKay, R. S., Johnson, S. & Sansom, B. How directed is a directed network? R. Soc. Open Sci. 7, 201138 (2020).

25. Kringelbach, M. L., Sanz Perl, Y., Tagliazucchi, E. & Deco, G. Toward naturalistic neuroscience: mechanisms underlying the flattening of brain hierarchy in movie-watching compared to rest and task. Sci. Adv. 9, eade6049 (2023).

26. Chopra, S. et al. The Transdiagnostic Connectome Project: an open dataset for studying brain-behavior relationships in psychiatry. Sci. Data 12, 923 (2025).

27. Cocuzza, C. V. et al. Brain network dynamics reflect psychiatric illness status and transdiagnostic symptom profiles across health and disease. Nat. Commun. 17, 6678 (2026).

28. Snaith, R. P. et al. A scale for the assessment of hedonic tone the Snaith–Hamilton Pleasure Scale. Br. J. Psychiatry 167, 99–103 (1995).

29. Gard, D. E., Gard, M. G., Kring, A. M. & John, O. P. Anticipatory and consummatory components of the experience of pleasure: a scale development study. J. Res. Pers. 40, 1086–1102 (2006).

30. Lovibond, P. F. & Lovibond, S. H. The structure of negative emotional states: comparison of the Depression Anxiety Stress Scales (DASS) with the Beck Depression and Anxiety Inventories. Behav. Res. Ther. 33, 335–343 (1995).

31. Yeo, B. T. T. et al. The organization of the human cerebral cortex estimated by intrinsic functional connectivity. J. Neurophysiol. 106, 1125–1165 (2011).

32. Morris, L. S. et al. Fronto-striatal organization: defining functional and microstructural substrates of behavioural flexibility. Cortex 74, 118–133 (2016).

33. Pizzagalli, D. A. & Roberts, A. C. Prefrontal cortex and depression. Neuropsychopharmacology 47, 225–246 (2022).

34. Whitton, A. E., Treadway, M. T. & Pizzagalli, D. A. Reward processing dysfunction in major depression, bipolar disorder and schizophrenia. Curr. Opin. Psychiatry 28, 7–12 (2015).

35. Zhang, W.-N., Chang, S.-H., Guo, L.-Y., Zhang, K.-L. & Wang, J. The neural correlates of reward-related processing in major depressive disorder: a meta-analysis of functional magnetic resonance imaging studies. J. Affect. Disord. 151, 531–539 (2013).

36. Kringelbach, M. L. The human orbitofrontal cortex: linking reward to hedonic experience. Nat. Rev. Neurosci. 6, 691–702 (2005).

37. Kringelbach, M. L. & Rolls, E. T. The functional neuroanatomy of the human orbitofrontal cortex: evidence from neuroimaging and neuropsychology. Prog. Neurobiol. 72, 341–372 (2004).

38. Choi, E. Y., Yeo, B. T. T. & Buckner, R. L. The organization of the human striatum estimated by intrinsic functional connectivity. J. Neurophysiol. 108, 2242–2263 (2012).

39. Haber, S. N. Corticostriatal circuitry. Dialogues Clin. Neurosci. 18, 7–21 (2016).

40. Grace, A. A. Dysregulation of the dopamine system in the pathophysiology of schizophrenia and depression. Nat. Rev. Neurosci. 17, 524–532 (2016).

41. Nestler, E. J. & Carlezon, W. A., Jr. The mesolimbic dopamine reward circuit in depression. Biol. Psychiatry 59, 1151–1159 (2006).

42. Russo, S. J. & Nestler, E. J. The brain reward circuitry in mood disorders. Nat. Rev. Neurosci. 14, 609–625 (2013).

43. Schultz, W., Dayan, P. & Montague, P. R. A neural substrate of prediction and reward. Science 275, 1593–1599 (1997).

44. Kaiser, R. H., Andrews-Hanna, J. R., Wager, T. D. & Pizzagalli, D. A. Large-scale network dysfunction in major depressive disorder: a meta-analysis of resting-state functional connectivity. JAMA Psychiatry 72, 603–611 (2015).

45. Kringelbach, M. L. & Berridge, K. C. The affective core of emotion: linking pleasure, subjective well-being, and optimal metastability in the brain. Emot. Rev. 9, 191–199 (2017).

46. Berridge, K. C. & Kringelbach, M. L. Affective neuroscience of pleasure: reward in humans and animals. Psychopharmacology 199, 457–480 (2008).

47. Berridge, K. C. & Kringelbach, M. L. Neuroscience of affect: brain mechanisms of pleasure and displeasure. Curr. Opin. Neurobiol. 23, 294–303 (2013).

48. Berridge, K. C. & Robinson, T. E. What is the role of dopamine in reward: hedonic impact, reward learning, or incentive salience? Brain Res. Rev. 28, 309–369 (1998).

49. Kringelbach, M. L. & Berridge, K. C. Towards a functional neuroanatomy of pleasure and happiness. Trends Cogn. Sci. 13, 479–487 (2009).

50. Kringelbach, M. L. et al. The entangled loop: Emotion, interference and hierarchy in the architecture of consciousness. (2026).

51. Reid, A. T. et al. Advancing functional connectivity research from association to causation. Nat. Neurosci. 22, 1751–1760 (2019).

52. Murty, V. P. et al. Resting state networks distinguish human ventral tegmental area from substantia nigra. NeuroImage 100, 580–589 (2014).

53. Pauli, W. M., Nili, A. N. & Tyszka, J. M. A high-resolution probabilistic in vivo atlas of human subcortical brain nuclei. Sci. Data 5, 180063 (2018).

54. Schaefer, A. et al. Local-global parcellation of the human cerebral cortex from intrinsic functional connectivity MRI. Cereb. Cortex 28, 3095–3114 (2018).

55. Tian, Y., Margulies, D. S., Breakspear, M. & Zalesky, A. Topographic organization of the human subcortex unveiled with functional connectivity gradients. Nat. Neurosci. 23, 1421–1432 (2020).

56. Horn, A. et al. Lead-DBS v2: towards a comprehensive pipeline for deep brain stimulation imaging. NeuroImage 184, 293–316 (2019).

57. Van Essen, D. C. et al. The WU-Minn Human Connectome Project: an overview. NeuroImage 80, 62–79 (2013).

58. Ashburner, J. A fast diffeomorphic image registration algorithm. NeuroImage 38, 95–113 (2007).

59. Cabral, J. et al. Metastable oscillatory modes emerge from synchronization in the brain spacetime connectome. Commun. Phys. 5, 184 (2022).

