## Supplementary Information for "Diminished ascending drive in transdiagnostic anhedonia shifts the cortico-subcortical hierarchy away from bottom-up dominance"

#### Contents


- Supplementary Figure 1. Network-level trophic levels across the seven Yeo cortical networks and the subcortex
- Supplementary Table 1. Reward-circuit ROI trophic levels across the three regression models
- Supplementary Table 2. Directed frontostriatal edge flow asymmetry across the three regression models
- Supplementary Table 3. Flow-asymmetry correlations with the Temporal Experience of Pleasure Scale
- Supplementary Discussion. Functional-connectivity control analysis
- Supplementary Table 4. Functional-connectivity families tested in the control analysis
- Supplementary Figure 2. Functional-connectivity control analysis

- Supplementary Table 5. Split-half reliability and within-half replication of the main measures

---

### Supplementary Figure 1

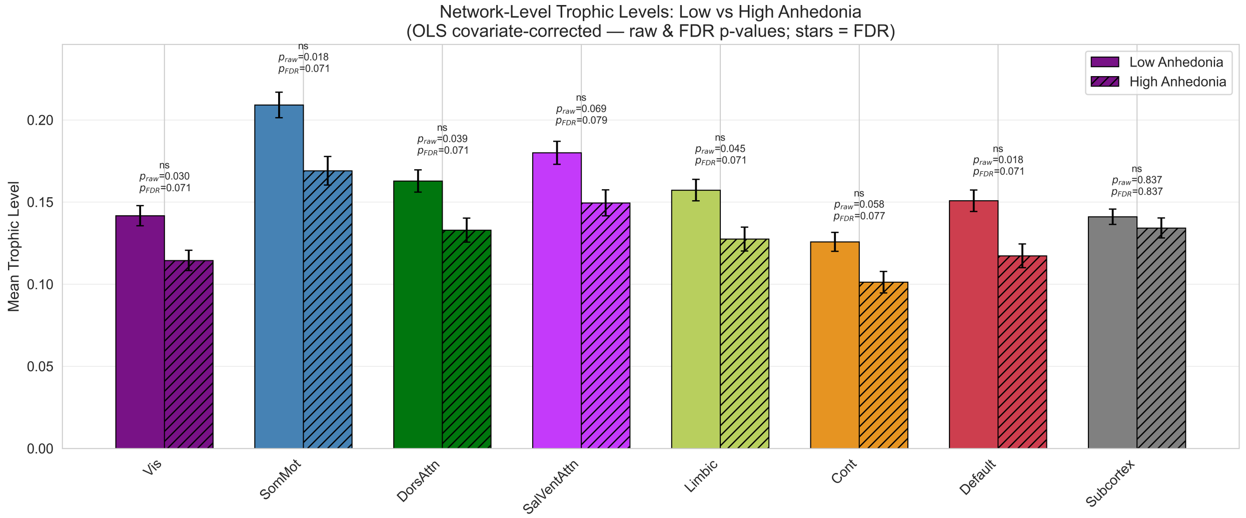


**Network-level trophic levels in low and high anhedonia.** Mean trophic level within each of the seven Yeo cortical networks and within the subcortex, compared between the low- and high-anhedonia groups. Bars give the group mean and error bars the standard error of the mean. Above each pair, $p_{\text{raw}}$ is the uncorrected p value from the covariate-corrected OLS model (anhedonia group, age, sex, patient/healthy-control status; N = 210) and $p_{\text{FDR}}$ the value after Benjamini–Hochberg correction across the eight comparisons. All seven cortical networks are lower in high anhedonia relative to each participant's most upstream region, with the largest differences in somatomotor and default-mode cortex, whereas the subcortex does not differ. No single network survives correction. Because levels are anchored to each participant's most upstream region, these single-network differences depend on that reference (see Methods in the main text). Vis, visual; SomMot, somatomotor; DorsAttn, dorsal attention; SalVentAttn, salience/ventral attention; Cont, frontoparietal control; ns, not significant after correction.

---

#### Supplementary Table 1

**Reward-circuit ROI trophic levels across the three regression models.**

| Reward ROI | Full β | Full p | Full q | Pat. β | Pat. p | Pat. q | +Dx β | +Dx p | +Dx q |
|---|---|---|---|---|---|---|---|---|---|
| NAc | −0.0089 | 0.289 | 0.405 | −0.0127 | 0.211 | 0.295 | −0.0135 | 0.187 | 0.262 |
| Caudate | +0.0035 | 0.665 | 0.665 | +0.0017 | 0.856 | 0.856 | +0.0013 | 0.896 | 0.896 |
| Putamen | +0.0052 | 0.605 | 0.665 | +0.0032 | 0.789 | 0.856 | +0.0031 | 0.797 | 0.896 |
| OFC | −0.0250 | 0.022 | 0.062 | −0.0287 | 0.023 | 0.057 | −0.0303 | 0.017 | **0.045** |
| mPFC/vmPFC | −0.0271 | 0.019 | 0.062 | −0.0328 | 0.014 | 0.057 | −0.0344 | 0.010 | **0.045** |
| ACC | −0.0232 | 0.026 | 0.062 | −0.0283 | 0.024 | 0.057 | −0.0295 | 0.019 | **0.045** |
| dlPFC | −0.0213 | 0.047 | 0.081 | −0.0260 | 0.034 | 0.059 | −0.0274 | 0.026 | **0.045** |

Group coefficients (high versus low anhedonia) for the mean trophic level of each of the seven reward regions of interest, under the three regression models reported throughout the paper. Full, the full sample adjusted for age, sex and patient/healthy-control status (N = 210); Pat., patients only with healthy controls excluded; +Dx, patients only with a broad three-level diagnostic category added. Negative coefficients denote a lower trophic level in high anhedonia, that is, a position closer to each participant's most upstream region. p is the uncorrected p value and q the Benjamini–Hochberg FDR-adjusted value across the seven ROIs within each model; values surviving $q < 0.05$ are in bold. All four cortical ROIs are significant uncorrected in every model and survive correction once broad diagnostic category is included, whereas the three striatal ROIs are non-significant throughout. NAc, nucleus accumbens; OFC, orbitofrontal cortex; mPFC/vmPFC, medial/ventromedial prefrontal cortex; ACC, anterior cingulate cortex; dlPFC, dorsolateral prefrontal cortex.

---

#### Supplementary Table 2

**Directed frontostriatal edge flow asymmetry across the three regression models.**

| Directed edge | Full β | Full p | Full q | Pat. β | Pat. p | Pat. q | +Dx β | +Dx p | +Dx q |
|---|---|---|---|---|---|---|---|---|---|
| dlPFC→NAc | +0.0110 | 0.0008 | **0.005** | +0.0101 | 0.004 | **0.026** | +0.0099 | 0.006 | **0.034** |
| vmPFC→NAc | +0.0087 | 0.014 | **0.027** | +0.0090 | 0.024 | 0.071 | +0.0094 | 0.018 | 0.052 |
| ACC→aPUT | +0.0490 | 0.009 | **0.027** | +0.0383 | 0.144 | 0.215 | +0.0434 | 0.091 | 0.136 |
| vmPFC→aPUT | +0.0117 | 0.047 | 0.071 | +0.0134 | 0.039 | 0.078 | +0.0140 | 0.026 | 0.052 |
| OFC→NAc | +0.0019 | 0.080 | 0.096 | +0.0007 | 0.596 | 0.596 | +0.0009 | 0.504 | 0.504 |
| ACC→NAc | −0.0125 | 0.456 | 0.456 | +0.0154 | 0.410 | 0.492 | +0.0163 | 0.386 | 0.463 |

Group coefficients (high versus low anhedonia) for the normalized flow-asymmetry index of each of the six theory-defined directed prefrontal-to-striatal edges, under the three regression models. Models are defined as in Supplementary Table 1 ($N = 210$ for the full sample). Positive coefficients denote a shift in the top-down direction in high anhedonia. p is the uncorrected p value and q the Benjamini–Hochberg FDR-adjusted value across the six edges within each model;

values surviving q < 0.05 are in bold. Rows are ordered by the strength of the full-model effect. The dlPFC→NAc edge is the only one to survive correction in all three models. aPUT, anterior putamen; other abbreviations as in Supplementary Table 1.

#### Supplementary Table 3

**Flow-asymmetry correlations with the Temporal Experience of Pleasure Scale.**

| Flow-asymmetry measure | TEPS scale | Pearson r | p | Spearman ρ | p | q |
|---|---|---|---|---|---|---|
| Cortex→subcortex | Total | −0.144 | 0.036 | −0.166 | 0.016 | 0.091 |
| PFC→striatum | Total | −0.159 | 0.021 | −0.196 | 0.004 | 0.091 |
| dlPFC→NAc | Total | −0.078 | 0.259 | −0.074 | 0.287 | 0.342 |
| ACC→aPUT | Total | −0.076 | 0.274 | −0.155 | 0.026 | 0.342 |
| vmPFC→NAc | Total | −0.044 | 0.523 | −0.044 | 0.526 | 0.523 |
| Cortex→subcortex | Consummatory | −0.139 | 0.044 | −0.157 | 0.023 | 0.221 |
| PFC→striatum | Consummatory | −0.160 | 0.020 | −0.181 | 0.009 | 0.201 |
| dlPFC→NAc | Consummatory | −0.089 | 0.200 | −0.046 | 0.512 | 0.400 |
| ACC→aPUT | Consummatory | −0.068 | 0.331 | −0.095 | 0.171 | 0.507 |
| vmPFC→NAc | Consummatory | −0.025 | 0.724 | −0.001 | 0.987 | 0.724 |
| Cortex→subcortex | Anticipatory | −0.113 | 0.103 | −0.116 | 0.093 | 0.257 |
| PFC→striatum | Anticipatory | −0.119 | 0.086 | −0.134 | 0.052 | 0.257 |
| dlPFC→NAc | Anticipatory | −0.050 | 0.471 | −0.061 | 0.376 | 0.528 |

| Flow-asymmetry measure | TEPS scale | Pearson r | p | Spearman ρ | p | q |
|---|---|---|---|---|---|---|
| ACC→aPUT | Anticipatory | −0.064 | 0.355 | −0.128 | 0.064 | 0.507 |
| vmPFC→NAc | Anticipatory | −0.050 | 0.475 | −0.043 | 0.538 | 0.528 |

Correlations between each subject's normalized flow-asymmetry index and the TEPS across the pooled sample (both anhedonia groups; N = 210). Because the TEPS is scored so that higher values denote greater hedonic capacity, a negative correlation indicates the same severity relationship as the positive SHAPS correlation reported in the main text. Every association is negative and therefore convergent in sign with the categorical group contrast. q is the Benjamini–Hochberg FDR-adjusted Pearson p value within the hypothesis-defined family, the secondary family for TEPS total (five tests) and the specificity family for the two subscales (ten tests). Pearson r is the corrected read-out and Spearman ρ the sensitivity check. No association survives correction, so the TEPS results are reported as a same-direction but sub-threshold trend. Within each scale, rows are ordered by the strength of the Pearson association. Abbreviations as in Supplementary Tables 1 and 2.

---

### Supplementary Discussion

##### *Functional-connectivity control analysis*

To establish that the directed-hierarchy findings reported in the main text are not recoverable from undirected connectivity, we repeated the key group analyses using static functional connectivity

(FC, the Fisher-z–transformed Pearson correlation between the parcellated regional time series) in place of the GEC-derived measures, under the identical covariate-corrected OLS design (age, sex, patient/healthy-control status; N = 210) and the same three-model and FDR conventions. Because FC is symmetric and edge-based rather than directed and nodal, each GEC measure was mapped to its closest undirected analogue. A region's or ROI's *trophic level*, a nodal and directional quantity, was matched by its **nodal FC strength**, the mean FC to all other regions, and each directional *flow asymmetry* across a boundary (cortex↔subcortex, PFC↔striatum) or along a directed edge was matched by the corresponding **mean undirected FC** across that boundary or between that ROI pair, a single value with no top-down or bottom-up arms to decompose. The same six families tested for trophic levels and flow asymmetry were re-run on the FC matrices (**Supplementary Fig. 2**).

**No FC metric reached significance in the covariate-adjusted model, and none survived FDR correction in any family** (**Supplementary Table 4**), in marked contrast to the directed-hierarchy measures, where the cortex − subcortex and PFC − striatum gaps, both boundary flow asymmetries, and the dlPFC→NAc edge were all robust and reproducible. FC was nonetheless not uniformly blank. At the compartment boundaries it carried a weak, same-direction trend consistent with the overall reduction in inter-compartmental coupling that the directed decomposition resolved into a significant effect. The directed measures are therefore best read as substantially more sensitive to the same reorganization, rather than as detecting an effect wholly absent from FC.

**Compartment FC.** No within- or cross-compartment FC difference reached significance, but the coefficients were consistently negative, indicating weaker coupling in high anhedonia in the same direction as the directed reduction. This held within cortex (β = −0.0158, $p = 0.38$), within subcortex (β = −0.0159, $p = 0.26$), and at the cortex↔subcortex boundary (β = −0.0165, 95% CI [−0.0406, +0.0076], $p = 0.18$). This cross-boundary trend mirrors, in undirected form, the overall reduction in inter-compartmental coupling that the directed two-arm decomposition resolved into a significant effect. What FC did *not* reproduce was the hierarchical *gap* effect. Because cortical and subcortical strength shifted together, the cortex − subcortex strength "gap", the FC analogue of the directed hierarchical gap that was the single strongest trophic effect ($p < 0.001$), was essentially zero (β = +0.0006, $p = 0.95$). The pattern held across the patients-only and diagnosis-adjusted models.

**Regional and network FC strength.** In the per-region analysis across all 232 regions, only six regions reached an uncorrected $p < 0.05$, fewer than the roughly twelve expected by chance at $\alpha = 0.05$, and **none survived FDR correction (0/232)**, with no Manhattan-plot structure resembling the 29-region cortical cluster seen for trophic levels. At the intermediate scale, none of the seven Yeo networks or the subcortex reached even uncorrected significance (all $p \geq 0.20$; all $q \approx 0.50$), and although every cortical network carried a negative coefficient, indicating lower whole-brain FC strength in high anhedonia, the effects were far too weak to interpret.

**Reward circuit.** None of the seven reward ROIs differed in nodal FC strength (all $p \geq 0.26$; all $q \geq 0.66$), and no cortical-versus-striatal contrast emerged. Collapsed to compartments, the PFC↔striatum boundary FC was again negative, in the same direction as the cortico-subcortical boundary and the directed frontostriatal coupling reduction, but a much fainter trend (β = −0.0077, 95% CI [−0.0366, +0.0212], $p = 0.60$). Within-PFC (β = −0.0114, $p = 0.48$), within-striatum (β = +0.0002, $p = 0.99$), and the PFC − striatum strength gap (β = +0.0008, $p = 0.95$) showed no difference, as at the whole-brain level. At finer grain the correspondence broke down. The six theory-defined frontostriatal edges were mixed in sign rather than uniformly weaker, and the dlPFC↔NAc edge, the strongest edge in the directed analysis, was flat in FC ($\beta = -0.0002$, $p = 0.99$); none of the six edges differed between groups (all $p \geq 0.41$; all $q \geq 0.95$). These edge- and NAc-level nulls, if anything, sharpen the specificity of the directed edge effect.

**Dimensional severity.** For completeness we also correlated the FC boundary and edge measures with SHAPS total across the pooled sample, uncorrected, as for the directed-flow severity analysis. Most were non-significant, but two reached an uncorrected threshold, the cortex↔subcortex boundary FC (Pearson $r = -0.18$, $p = 0.010$; Spearman $\rho = -0.15$, $p = 0.029$) and the vmPFC↔anterior-putamen edge ($r = -0.14$, $p = 0.043$; $\rho = -0.22$, $p = 0.002$), both *negative*, meaning that weaker undirected coupling accompanied higher anhedonia. These are weak, uncorrected associations on measures whose primary group contrasts were null, so, unlike the directed-flow asymmetries, which were significant at both the group and dimensional levels, they do not constitute a reliable FC signal and are reported only for transparency.

Taken together, the functional-connectivity control reproduces the directed-hierarchy signature only weakly and partially. FC retained a same-direction but sub-significant trend toward reduced cross-boundary coupling, cortico-subcortical and more faintly frontostriatal, consistent with the overall drop in inter-compartmental coupling that the directed decomposition revealed. It did not reproduce the *directional* reorganization, neither the hierarchical gaps and their flow-asymmetry counterparts nor the dlPFC→NAc edge, all of which carried the significant, reproducible effects. The anhedonia-related reorganization is therefore expressed primarily in the *direction* of information flow and in regions' *hierarchical position*, and only faintly in the symmetric coupling magnitude that FC measures. This is best read as a difference in sensitivity rather than a categorical presence or absence. The directed, generative-model read-outs resolve into robust effects a reorganization that undirected connectivity registers, at most, as a weak trend, which is exactly what motivates the directed approach over conventional functional connectivity.

---

#### Supplementary Table 4

**Functional-connectivity families tested in the control analysis.**

| FC family (undirected analogue) | Metrics tested | Uncorrected $p < 0.05$ | Survive FDR |
|---|---|---|---|
| Compartment FC (cortex, subcortex, boundary and nodal strength) | 6 | 0 | — (reported uncorrected) |
| Regional FC strength | 232 | 6 | 0 |
| Network FC strength (Yeo-7 and subcortex) | 8 | 0 | 0 |
| Reward-circuit ROI FC strength | 7 | 0 | 0 |

| FC family (undirected analogue) | Metrics tested | Uncorrected $p < 0.05$ | Survive FDR |
| --- | --- | --- | --- |
| Reward compartment FC (PFC, striatum, boundary and nodal strength) | 6 | 0 | — (reported uncorrected) |
| Frontostriatal reward edges | 6 | 0 | 0 |

Number of metrics tested in each of the six functional-connectivity families, the number reaching an uncorrected $p < 0.05$, and the number surviving Benjamini–Hochberg FDR correction within that family. All models are the covariate-corrected OLS design used for the directed measures ($N = 210$). No family yields a single FDR-surviving metric. The two compartment families are reported uncorrected because they comprise a small, pre-specified set rather than a screen.

### Supplementary Figure 2

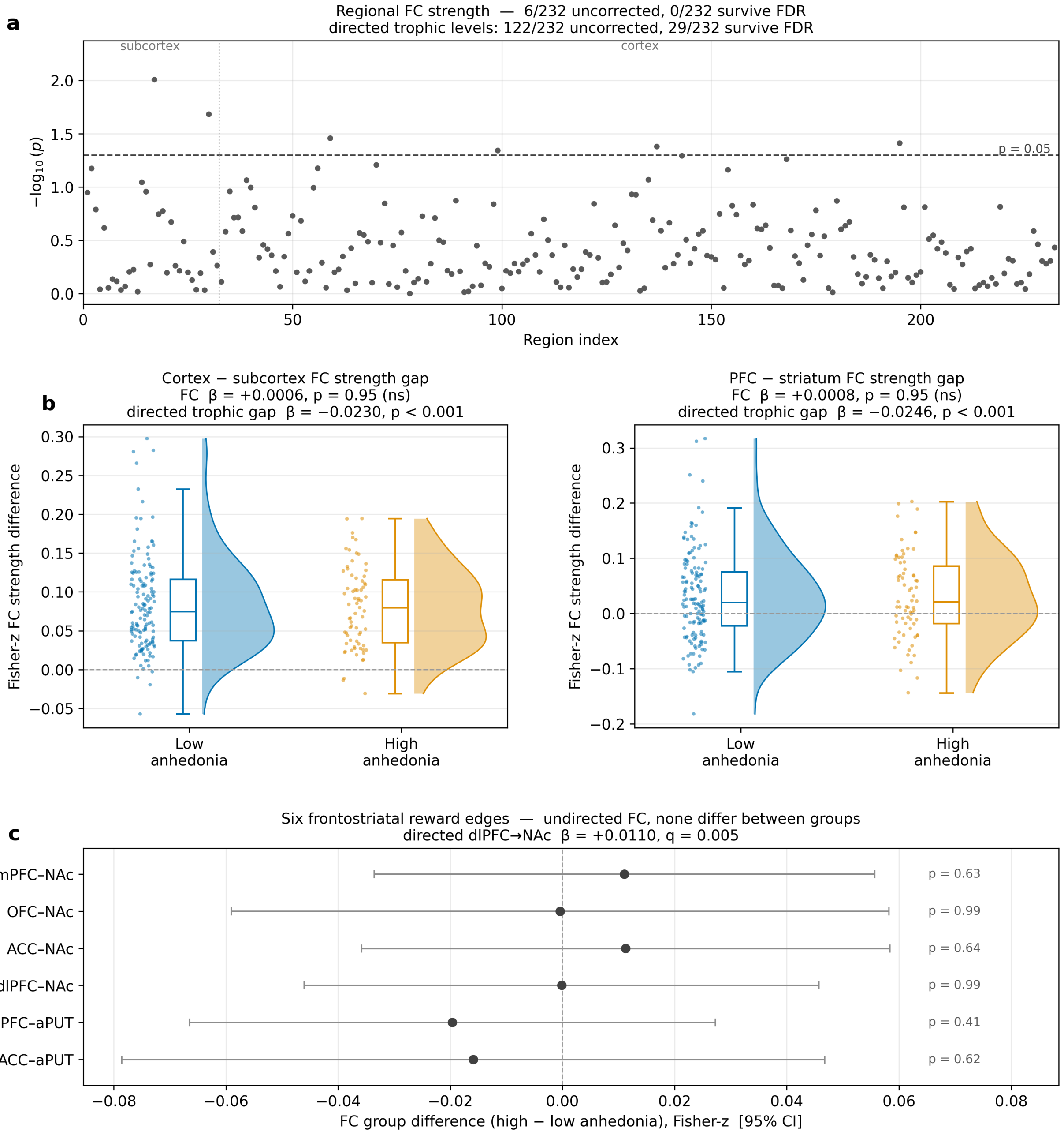


**The directed-hierarchy group analyses are not reproduced by undirected functional connectivity.** The key group analyses repeated on undirected FC under the covariate-corrected OLS design used for the directed measures (age, sex, patient/healthy-control status; N = 210). Each panel is annotated with the matched directed result for contrast. **(a)** Per-region FC nodal strength across all 232 regions, plotted as −log10(p) against region index, with the dashed line marking the uncorrected p = 0.05 threshold and the dotted vertical line separating subcortex from cortex. No region survives FDR correction, against 29 for the directed trophic levels of Fig. 3a of the main text. **(b)** The FC analogues of the cortex − subcortex and PFC − striatum hierarchical gaps, whose group difference is essentially zero, against the directed gaps of Figs 3e and 4b of the main text, which fall from positive to about zero. Each panel shows a half violin (kernel density, normalized to equal width), a box (median and interquartile range, whiskers 1.5 × interquartile range, outliers omitted) and one dot per participant. **(c)** Group difference in undirected FC for each of the six frontostriatal reward edges, with dots giving the coefficient and bars the 95% confidence interval. Every interval crosses zero, against the directed edge results of Supplementary Table 2. aPUT, anterior putamen; other abbreviations as in Supplementary Table 1.

---

#### Supplementary Table 5

**Split-half reliability and within-half replication of the main measures.**

| Measure | r | SB | ICC | Half 1 β (p) | Half 2 β (p) | Full β (p) |
|---|---|---|---|---|---|---|
| Cortex–subcortex gap | 0.66 | 0.80 | 0.66 | −0.021 (<0.001) | −0.027 (<0.001) | −0.023 (<0.001) |
| Cortex–subcortex flow | 0.71 | 0.83 | 0.70 | +0.015 (<0.001) | +0.018 (<0.001) | +0.015 (<0.001) |

| Measure | r | SB | ICC | Half 1 β (p) | Half 2 β (p) | Full β (p) |
|---|---|---|---|---|---|---|
| PFC–striatum gap | 0.62 | 0.76 | 0.61 | −0.022 (0.004) | −0.032 (<0.001) | −0.025 (<0.001) |
| PFC–striatum flow | 0.69 | 0.82 | 0.68 | +0.017 (<0.001) | +0.019 (<0.001) | +0.016 (<0.001) |
| Trophic coherence | 0.67 | 0.80 | 0.62 | −0.000 (0.98) | +0.000 (0.77) | +0.000 (0.69) |
| vmPFC→NAc | 0.41 | 0.58 | 0.38 | +0.006 (0.09) | +0.015 (0.006) | +0.009 (0.01) |
| vmPFC→aPUT | 0.51 | 0.68 | 0.49 | +0.013 (0.05) | +0.019 (0.02) | +0.012 (0.05) |
| dlPFC→NAc | 0.34 | 0.51 | 0.34 | +0.008 (0.09) | +0.016 (0.001) | +0.011 (<0.001) |
| OFC→NAc | 0.24 | 0.39 | 0.23 | – | – | – |

Split-half reliability and within-half replication of the main measures. GECs were refitted separately on half 1 (run 1, AP + PA) and half 2 (run 2, AP + PA). r, Pearson correlation between halves (N = 210); SB, Spearman–Brown-corrected reliability of the full four-run measure estimated from r; ICC, absolute-agreement intraclass correlation ICC(A,1) for a single half; flow, normalized flow-asymmetry index. Group coefficients (high versus low anhedonia) with uncorrected p values are from the primary covariate model (age, sex, patient/healthy-control status; N = 206), fitted separately within each half and on the full data. –, not tested within halves (not significant in the full data). The ACC→NAc and ACC→anterior-putamen edges could not be assessed, because in some participants both directions carried zero weight in at least one half, leaving the normalized index undefined. aPUT, anterior putamen; other abbreviations as in

Supplementary Table 1.
